# Role of Astrophorina sponges (Demospongiae) in food-web interactions at the Flemish Cap (NW Atlantic)

**DOI:** 10.1101/2023.05.06.539722

**Authors:** Tanja Stratmann, Francisco Javier Murillo, Mar Sacau, Mariano Koen Alonso, Ellen Kenchington

## Abstract

Deep-sea sponges are important contributors to carbon and nitrogen cycling due to their large filtration capacity. Species of the suborder Astrophorina form dense sponge grounds in the North Atlantic, where they serve as prey for spongivores, but also have non-trophic interactions with commensal epi- and endobionts. At the Flemish Cap (NW Atlantic) Astrophorina sponges are present in four previously described deep-sea epifaunal assemblages: the deep-sea coral assemblage, the lower slope assemblages 1 and 2, and the deep-sea sponge assemblage. To investigate their role in trophic and non-trophic interactions at the Flemish Cap, we developed trophic-/ non-trophic interaction web models for each of the four faunal assemblages using the published literature. By excluding the sponges from the models, we estimated how many trophic, facultative and obligatory non-trophic links would be lost, and how this removal affected food-web properties (number of compartments, links, link density, and connectance). Astrophorina sponges were mostly linked via facultative non-trophic links to 60, 59, 86, and 92 compartments in the deep-sea coral, the lower slope 1 and 2, and the deep-sea sponge assemblages, respectively. Direct trophic links only existed to Echinasteridae and Pterasteridae. As removing Astrophorina sponges from the interaction webs of the different assemblages had the highest impact on food-web properties compared to removing any other fauna present, these sponges were considered “highest impact taxa”. They were also identified, along with sea pens, as “structural species”/ “habitat formers” and “foundation species” based on non-trophic interactions in the deep-sea coral and deep-sea sponge assemblages.

## 1. INTRODUCTION

Sponges (Porifera) are important filter feeders that efficiently filter large amounts of bacterioplankton, dissolved organic matter (DOM), and particulate organic matter (POM) out of the water column (Bart et al. 2020, 2021b c, Maier et al. 2020). For example, sponge grounds on the continental shelf of northern Norway pump 250 million m^3^ of water every day consuming 50 t C d^-1^ (Kutti et al. 2013) and 1 m^2^ sponge ground at the Karasik Seamount (Langseth Ridge, central Arctic Ocean) processes all the water in the overlying 600 m of water column in one year (Morganti et al. 2022). Sponge grounds consisting of the demosponge *Geodia* sp. are also very important for nitrogen cycling. Hoffmann et al. (2009) estimated that sponge-mediated nitrification rates transform up to 16 mmol N m^-2^ d^-1^ and remove 2.7 mmol N m^-2^ d^-1^ as N_2_ which implies that nitrogen removal rates could be 2 to 10 times higher than nitrogen removal rates at continental slopes (Middelburg et al. 1996, Seitzinger & Giblin 1996, Hoffmann et al. 2009).

The biomass in sponge grounds consisting of the demosponge suborder Astrophorina (primarily species of *Geodia* sp., *Thenea* sp., and *Stryphnus* sp.) is not only controlled by the availability of food sources in the water column (i.e., bottom-up control), but also by the presence of spongivores/ predators (i.e., top-down control). Bart et al. (2021a) described a deep-sea sponge loop in which brittle stars (Ophiuroidea) either consume detritus produced by *Geodia barretti* which was previously fed with DOM and POM, a process the authors called “deep-sea detrital sponge loop” or by direct predation of the brittle stars on the DOM- and POM-fed *Geodia barretti* which is called the “deep-sea predatory sponge loop”. In both cases, sponges recycle DOM and make it available as food source to higher trophic levels.

Besides these trophic links, sponges are also involved in non-trophic links by providing three dimensional habitats for commensal epi- and endobionts. For example, a study of seven sponge species collected at the US-Atlantic continental shelf and slope revealed a mean density of (mean±SD) 1976±3267 ind^-1^ associated fauna l^-1^ sponge (Fiore & Cox Jutte 2010). For comparison, tunicates had a mean density of associated fauna of 642±619 ind^-1^ associated fauna l^-1^ tunicate volume (Fiore & Cox Jutte 2010). Most of the fauna associated with sponges were polychaetes (93%), whereas tunicates hosted polychaetes (28%), decapods (28%), and amphipods (32%) (Fiore & Cox Jutte 2010). Even when only larger invertebrate megabenthos, which is visible on photos, is considered, sponge grounds host a higher diversity and density of benthos than adjacent areas without sponges (Beazley et al. 2013). In the Flemish Cap on the Canadian continental margin of the Atlantic Ocean, non-sponge grounds had a mean megabenthos density of (mean±SE) 33±3 ind m^-1^ and a mean Shannon diversity *H’* index of 1.3±0.04, whereas sponge grounds had a mean megabenthos density of 65±5 ind m^-1^ and mean *H’* index of 2.1±0.02 (Beazley et al. 2013). Particularly the demersal fish community benefits from the presence of sponge grounds and/ or dense coral gardens with 0.106 ind. m^-1^ observed in areas without corals or sponge grounds, 0.111 ind. m^-1^ in areas with dense sponge cover, 0.187 ind. m^-1^ in areas with dense coral cover, and 0.131 ind. m^-1^ in areas with a mix of sponge grounds and coral gardens (Devine et al. 2020).

Traditionally, the study of trophic links is based on topological food webs (e.g., Hanz et al., 2022; Morato et al., 2016) and energy flow webs (e.g., Soetaert & van Oevelen 2009, van Oevelen et al. 2009, Stratmann et al. 2018, de Jonge et al. 2020, Stratmann 2023), whereas non-trophic links have been studied via non-trophic or interaction webs (e.g., Salinas et al., 2023). In 2015, van der Zee et al. (2016) introduced a modeling approach to combine the assessment of trophic- and non-trophic links in interaction webs. Stratmann et al. (2021) developed this approach further to assess the role of polymetallic nodules for food-web integrity in abyssal plains of the Southeast and Central Pacific Ocean. In that study, the authors identified stalked sponges that grow attached to the nodules as so-called “highest impact taxa”, i.e., taxa whose removal resulted in the largest changes in food-web properties, such as number of species or interaction-web links.

Here, we used the models from Stratmann et al. (2021) to (1) investigate the role of sponges of the suborder Astrophorina for trophic and non-trophic interactions in four different deep-sea epifaunal assemblages (1 coral assemblage, 2 lower continental slope assemblages, 1 sponge assemblage) at the Flemish Cap, and to (2) identify the highest impact taxon of each assemblage.

## 2. MATERIALS & METHODS

### 2.1 Study site

The Flemish Cap is a bank in the high seas of the continental margin off Newfoundland, Canada, with a radius of ∼200 km at the 500 m isobath and minimum depth of approximately 122 m (Fig. 1). It is considered both a bioregion and an ecosystem production unit, based on analyses of a suite of physiographic, oceanographic, and biotic variables (NAFO 2014). It is treated as a discrete unit, Northwest Atlantic Fisheries Organization (NAFO) Division 3M, for the management of commercial fisheries prosecuted on the bank, and for ecosystem summaries informing fisheries management. There are steep slopes to the east and south, below 1,000 m depth, but more gradual gradients to the north and west. It is separated from the Grand Banks by the Flemish Pass, a 1,200 m deep, mid slope channel. Two major ocean currents influence this area: the Labrador Current, flowing from the north, and the North Atlantic Current, which represents an extension of the warm Gulf Stream. When the Labrador Current reaches the Flemish Pass, it bifurcates with the major branch flowing southwards to the south eastern slope of the Grand Bank; meanwhile, the side branch circulates clockwise around the Flemish Cap.

**Figure 1.**
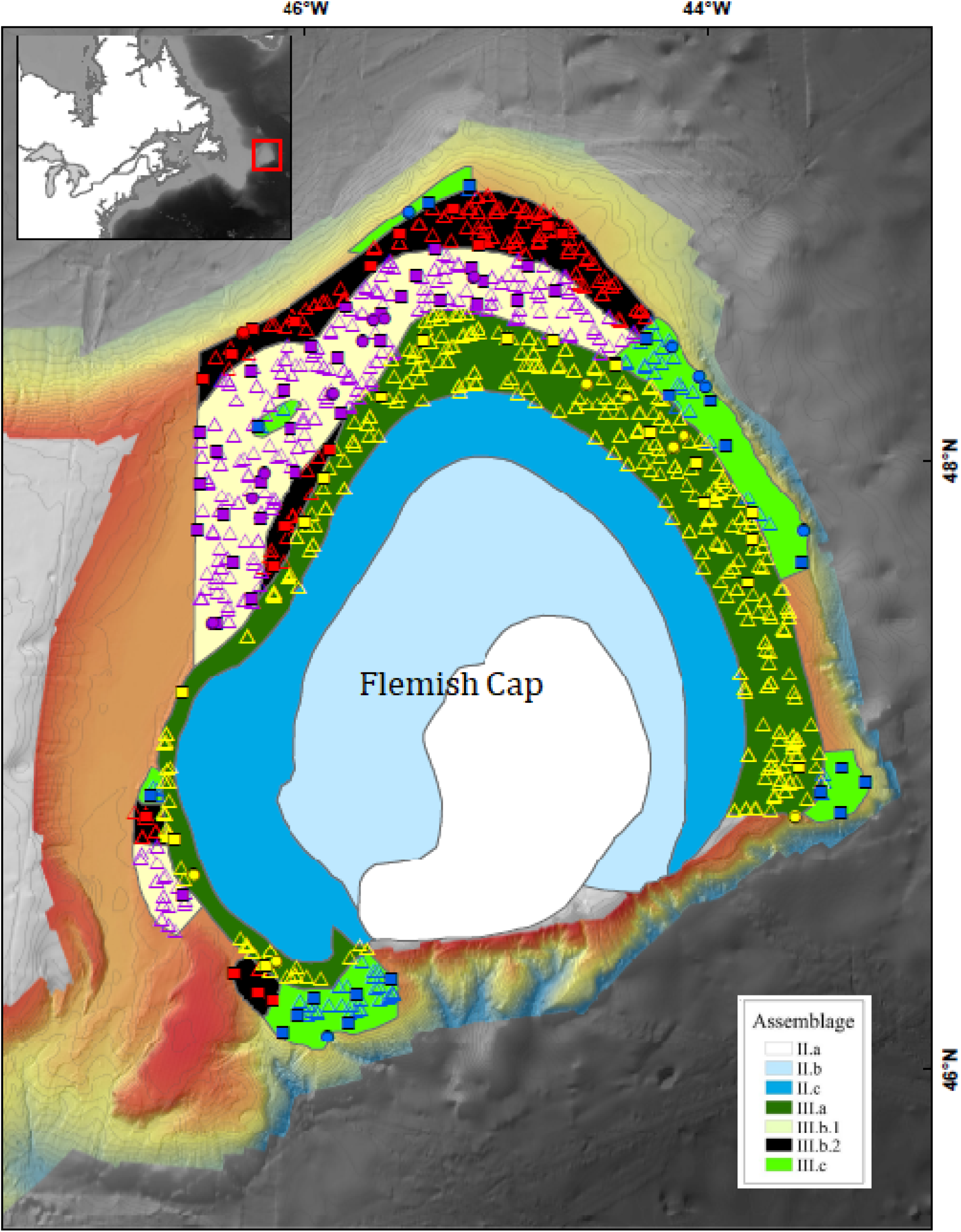
Study area on Flemish Cap, northwest Atlantic (inset), and data distributions overlain on the benthic assemblages delineated by Murillo et al. (2016). Open triangles represent research vessel survey trawl start positions, solid squares represent box core locations and solid circles represent rock/scallop dredge locations. Yellow indicates samples from the deep-sea coral assemblage, purple indicates samples from the lower slope assemblage 1, red indicates samples from the lower slope assemblage 2, blue represents samples from the deep-sea sponge assemblage. Multibeam bathymetry between 700 and 2000 m is shown in the background grading from shallow (reds) to deep (blues). The depth range of the samples are provided in Table 1.

**Table 1.** Summary of data collected with each of the three sampling gears in each of the four benthic community types described by Murillo et al. (2016).

| **Benthic community** | **No. Lofoten trawl sets** | **Depth range (m)** | **No. rock dredge sets** | **Depth range (m)** | **No. box cores** | **Depth range (m)** |
| --- | --- | --- | --- | --- | --- | --- |
| Deep-sea coral assemblage | 376 | 470–974 | 7 | 605–956 | 18 | 688–1125 |
| Lower slope assemblage 1 | 301 | 775–1202 | 8 | 935–1132 | 31 | 800–1181 |
| Lower slope assemblage 2 | 151 | 855–1440 | 1 | 1264–1294 | 17 | 981–1391 |
| Deep-sea sponge assemblage | 62 | 653–1355 | 6 | 1316–1589 | 19 | 750–1537 |

### 2.2 Sample collection and data compilation

Murillo et al. (2016) identified the invertebrate epibenthos from the catches of the 2007 EU Flemish Cap bottom-trawl survey. Statistical analyses of those data found twelve significantly different epibenthic assemblages with two of those shared between the Flemish Cap and the Tail of the Grand Bank in the deeper slope areas (Murillo et al. 2016). Four of those assemblages (Table 2) were used to delineate data sets for our study of food-web interactions on the Flemish Cap, as they overlapped with the distribution of macrobenthic data collected in this region. All are deep-sea assemblages from the continental slopes and include a deep-sea coral assemblage, two lower slope assemblages (lower slope 1, lower slope 2), and a deep-sea sponge assemblage (Murillo et al. 2016a) (Fig. 2).

**Figure 2.**
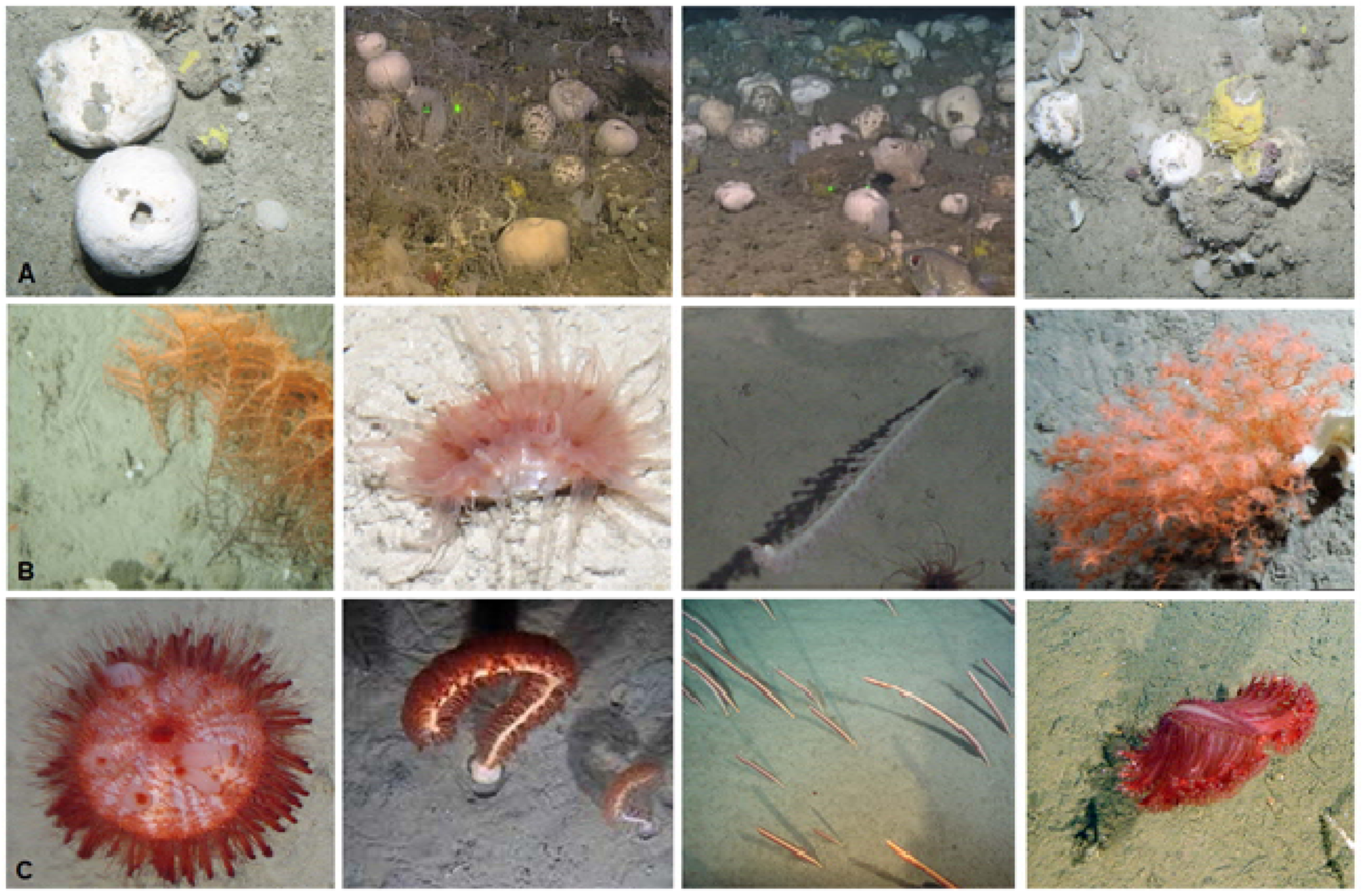
Fauna characteristic for the four benthic community types. **(A)** Deep-sea sponge assemblage typified by high biomass of large sponges mainly from the suborder Astrophorina; **(B)** Deep-sea coral assemblage showing left to right typical species: Black coral *Stauropathes arctica* (φ = 0.62), cup coral *Flabellum alabastrum* (φ = 0.59), sea pen cf. *Funiculina quadrangularis* (φ = 0.52), and small gorgonian coral *Acanella arbuscula* (φ = 0.41) [soft coral *Heteropolypus sol* (φ = 0.46) is not shown]; **(C)** Lower slope assemblages showing left to right typical species: Sea urchin *Phormosoma placenta* (φ = 0.55), sea pens *Anthoptilum grandiflorum* (φ = 0.45), *Balticina finmarchica* (φ = 0.41) and *Pennatula aculeata* (φ = 0.40) [sea stars *Bathybiaster vexillifer* (φ = 0.49) and *Zoroaster fulgens* (φ = 0.42), and *Funiculina quadrangularis* (φ = 0.49) are not shown].

**Table 2.** List and properties of benthic assemblages (following (Murillo et al. 2016a)) used to organize the analyses in this study. Φ, the Phi coefficient, is a measure of the strength of association of each species with the assemblages. Following (Chytrý & Tichý 2003), sharpness is defined as the number or quality of diagnostic species in a cluster group relative to its mean species richness. Uniqueness is maximum if none of the diagnostic species in a cluster are considered diagnostic in other cluster groups.

| Benthic assemblage | Representative taxa | Sharpness | Uniqueness | Mean taxon richness (SD) | Mean depth (SD) |
| --- | --- | --- | --- | --- | --- |
| Deep-sea coral assemblage | Black coral *Stauropathes arctica* ($\phi = 0.62$), the cup coral *Flabellum alabastrum* ($\phi = 0.59$), the sea pen *Funiculina quadrangularis* ($\phi = 0.52$), the deep-sea mushroom soft coral *Heteropolypus sol* ($\phi = 0.46$) and the small gorgonian coral *Acanella arbuscula* ($\phi = 0.41$). | 53.6 | 0.88 | 32 (15) | 658 (124) |
| Lower slope assemblage 1 | The sea urchin *Phormosoma placenta* ($\phi = 0.55$), the sea stars *Bathybiaster vexillifer* ($\phi = 0.49$) and *Zoroaster fulgens* ($\phi = 0.42$), and the sea pens *Funiculina quadrangularis* ($\phi = 0.49$), *Anthoptilum grandiflorum* ($\phi = 0.45$), *Balticina finmarchica* ($\phi = 0.41$) and *Pennatula aculeata* ($\phi = 0.40$). | 54.8 | 0.77 | 17 (8) | 1004 (134) |
| Lower slope assemblage 2 | Similar to lower slope assemblage 1, but with a wider bathymetric range. | 39.7 | 0.39 | 9 (5) | 1092 (208) |
| Deep-sea sponge assemblage | Typified by high biomass of large sponges mainly from the suborder Astrophorina. | 66.4 | 1.60 | 33 (10) | 1136 (190) |

Data used in this study were obtained from two sources. Fish and invertebrates were recorded from the catches of the 2011–2019 EU Flemish Cap bottom-trawl research surveys, conducted by the Instituto Español de Oceanografía together with the Instituto de Investigaciones Marinas and the Instituto Português do Mar e da Atmosfera. The surveys sampled the Flemish Cap and the eastern side of Flemish Pass between 470 and 1,440 m depth (Fig. 1), following a depth-stratified random sampling design. Surveys were conducted with standardized sets of a Lofoten bottom trawl (mean swept area of ≈39,000 m^2^ each) aboard the Spanish RV *Vizconde de Eza*. Samples were processed as described by Vázquez et al. (2014). Those data were complemented by data from the NEREIDA surveys (Durán Muñoz et al. 2012, NAFO 2013) undertaken aboard the Spanish RV *Miguel Oliver* in the Flemish Pass and Flemish Cap at depths between 605 and 1,589 m (Table 1, Figure 1). NEREIDA surveys were carried out during spring and summer of 2009 and 2010. Infaunal samples were collected using a mega boxcore (USNEL type, 0.25 m^2^ sampling area), and benthic macrofauna were processed as detailed in Barrio Froján et al. (2016). For hard seabed bottom types (compacted sands, gravels, and rock) a rock dredge/scallop gear was deployed, which consisted of a rectangular metal collar, coupled with a coarse mesh net protected by a rubber mat. The device was towed for approximately 1 km along the seabed allowing the rectangular metal mouth of the dredge to dig into the substrate, parts of which are then retained in the sample net. The towing speed was between 2 and 3 knots (NAFO 2013). All three data sets included positional information and the biomass (and abundance for some) of each species identified in every sample.

We used the benthic assemblage boundaries, qualitatively mapped by Murillo et al. (2016), to partition the data in each assemblage (Table 2), using the spatial join function in ArcMap version 10.7 (ESRI 2019). For each record, taxonomic position was assigned following the World Register of Marine Species consulted in 2021 (Horton et al. 2018). Literature searches were subsequently conducted for each taxon to support 1) designation of dominant adult feeding strategy, and diet items for elucidation of trophic links, 2) adult size class [meiobenthos >32 µm; macrobenthos >250 µm/ >500 µm; invertebrate megabenthos >1 cm], 3) non-trophic associations with Astrophorina sponges, 4) non-trophic associations with other taxa (see (Stratmann et al. 2023)).

### 2.3 Trophic/ non-trophic interaction web modeling

The interaction-web modelling exercise requires the constructions of trophic and non-trophic interaction web matrices (*TI* matrix and *NTI* matrix, respectively). To build these matrices, data was compiled in three arrays. Array 1 identifies records of all taxa that are present, array 2 lists their feeding types/ diet preferences, and array 3 includes information about any commensal relationships (i.e., relationships in which one species benefits while the other species is not affected) among taxa. These non-trophic links can be obligatory or facultative. Obligatory non-trophic links include, e.g., the dependence of deep-sea incirrate octopods on stalked sponges in the abyssal central Pacific where these octopods lay their eggs around the stalk of sponges and brood them (Purser et al. 2016). In comparison, a facultative non-trophic link exists for example when a sea anemone of the family Hormathiidae grows physically attached to Astrophorina sponges at the Langseth Ridge (central Arctic Ocean) (Stratmann et al. 2022) where the sponges serve as hard substrate. However, these sea anemones also grow on rocks or larger gravel (Stratmann, personal observation). Data from arrays 1 and 2 contribute to the development of the *TI* matrix, and data from arrays 1 and 3 are used to develop the *NTI* matrix.

The *TI* matrix links all faunal compartments of an assemblage to their prey/ food sources via all trophic links that have been reported in the literature. Hence, when a specific fish species preferably predates upon crustaceans, we linked this fish compartment via trophic links to all faunal compartments in the trophic interaction web that contained crustaceans. For species that were not predators, but e.g., bacterivores, filter and suspension feeders, or deposit feeders, we included extra food sources, such as bacteria, sedimentary detritus, or suspended detritus (Table 3). To achieve a square *TI* matrix with identical dimensions like the square *NTI* matrix, we furthermore added a compartment called “Astrophorina sponges” which is part of the *NTI* matrix due to the focus of this paper on the role of Astrophorina sponges, but it had no trophic links to any other compartment in the *TI* matrix. Rows in the matrix are prey, while columns are predators; the number 1 describes a trophic link and the number 0 means that there is no trophic link.

**Table 3.** List of extra food sources that were included in the trophic interaction matrices of the four benthic assemblages.

| Benthic assemblage | Extra food sources |
| --- | --- |
| Deep-sea coral assemblage | (Endosymbiotic chemosynthetic -) bacteria, carrion, dissolved organic matter, fish eggs and larvae, microbial mats, sedimentary detritus, sedimentary and suspended diatoms, suspended particulate organic matter |
| Lower slope assemblage 1 | (Endosymbiotic chemosynthetic -) bacteria, carrion, dissolved organic matter, fish eggs and larvae, microbial mats, (labile-) sediment detritus, sedimentary and suspended diatoms, suspended particulate organic matter, macro-/ megabenthic Foraminifera |
| Lower slope assemblage 2 | (Endosymbiotic chemosynthetic -) bacteria, carrion, dissolved organic matter, fish eggs and larvae, microbial mats, (labile-) sediment detritus, phytodetritus, suspended diatoms and particulate organic matter, macro-/ megabenthic Foraminifera |
| Deep-sea sponge assemblage | Bacteria, carrion, dissolved organic matter, macro-/ megabenthic Foraminifera, (labile-) sedimentary detritus, phytodetritus, suspended particulate organic matter, fish eggs |

The *NTI* matrix has the same dimensions as the *TI* matrix and includes all non-trophic links among faunal compartments, the Astrophorina sponge compartment, and other faunal compartments. The matrix is filled from rows or source to columns or utilizer, hence, a link from Astrophorina to Polychaeta implies that this specific polychaete has a non-trophic link to Astrophorina sponges. An obligatory non-trophic link is defined in the matrix as 1 and a facultative non-trophic link is defined as 2. We prepared both matrices with the functions *fillTImatrix* and *fillNTImatrix* in the Supplementary Material, which are modified versions of the same functions from Stratmann et al., (2021).

The role of a specific faunal compartment, or the Astrophorina compartment for trophic and non-trophic links in a specific assemblage, was assessed with the function *Remove* in the Supplementary Material and in Stratmann et al., (2021). This function identifies all trophic, facultative non-trophic, and obligatory non-trophic links a selected compartment has, as all of them will be lost during a “level 1 loss”. This level 1 loss functions as follows: A compartment A has a trophic or a non-trophic link to compartment B (or the Astrophorina compartment) so that removing compartment B will cause the immediate loss of compartment A. Hence, compartment A disappears in a level 1 loss. Subsequently, the function estimates so-called “maximum secondary losses”, i.e., it identifies all compartments that will be lost because they have a trophic, a facultative non-trophic, or an obligatory non-trophic link with a compartment that has been lost during the level 1 loss. The maximum secondary loss is not a theoretical secondary extinction in a traditional sense (see e.g., Brodie et al., 2014), but we assume that the key trophic or non-trophic link that allows the taxon to survive is the link that is associated with the compartment that was removed, regardless of how many other links exist between the taxon and other compartments. This approach assumes, therefore, no resilience for the impacted taxa that still have links. The function continues to iteratively assess any further “maximum higher-level losses” until no more compartments are lost, and the results are presented in a summary table (Supplementary Material).

### 2.4 Network indices

Based on the *TI* matrices, we calculated the network indices “number of interaction web compartments” *S*, “number of network links” *L*, “link density” *D* (D = L/S), and “connectance” *C* (C = L/ S^2^), which was the fraction of all realized links in comparison to all links that were possible (Pimm et al. 1991).

### 2.5 Assessment of the absence of specific faunal compartments on network indices

We studied the effects of removing Astrophorina sponges from the various faunal assemblages: how did this affect the network indices *S*, *L*, *D*, and *C*, which taxa were lost due to trophic links, which ones due to obligatory and facultative links, were these losses level 1 losses, maximum secondary losses, or maximum higher-level losses? Furthermore, we assessed the importance of representative taxa of the individual faunal assemblages on each assemblage by removing them from the *TI* and *NTI* matrices and by calculating changes in network indices.

We also identified which taxa had the most trophic interactions as prey (i.e., which taxa had most predators) and which taxa had most trophic interactions as predator (i.e., which taxa predated upon most prey) and how their removal altered the network indices.

In a last step, we iteratively removed each taxon from the *TI*/ *NTI* matrices and calculated the change in the network indices for each individual iteration. The taxon whose removal caused the highest absolute change in one or more of the network indices *S*, *L*, *D*, and *C* was considered a “highest impact taxon”.

## 3. RESULTS

### 3.1 Food webs of the different faunal assemblages at the Flemish Cap

#### 3.1.1 Deep-sea coral assemblage

The deep-sea coral assemblage consisted of 252 faunal interaction-web compartments including protozoan and metazoan meiobenthos (0.79%), macrobenthos (8.33%), invertebrate megabenthos (57.1%), and fish (33.7%). Most of the taxa were carnivores (64.3%) and filter/ suspension feeders (19.0%) (Fig. 3A) and the dominant phyla were Arthropoda (15.1% of all faunal compartments), Chordata (34.1%), Cnidaria (15.1%), Echinodermata (13.9%), and Mollusca (11.5%) (Fig. 4A). All compartments were linked via 10,211 trophic links with their respective food sources (Fig. 5, Table 3, Table 4) and the non-trophic interaction web comprised 248 links (Fig. 5). The link density was 39 and the connectance was 0.15 (Table 4). The predator with most trophic links in the deep-sea coral assemblage was the thorny skate (*Amblyraja radiata*) and the prey with most trophic links was krill of the order Euphausiacea. The taxon with most non-trophic (commensal) links was polychaetes of the family Polynoidae.

**Figure 3.**
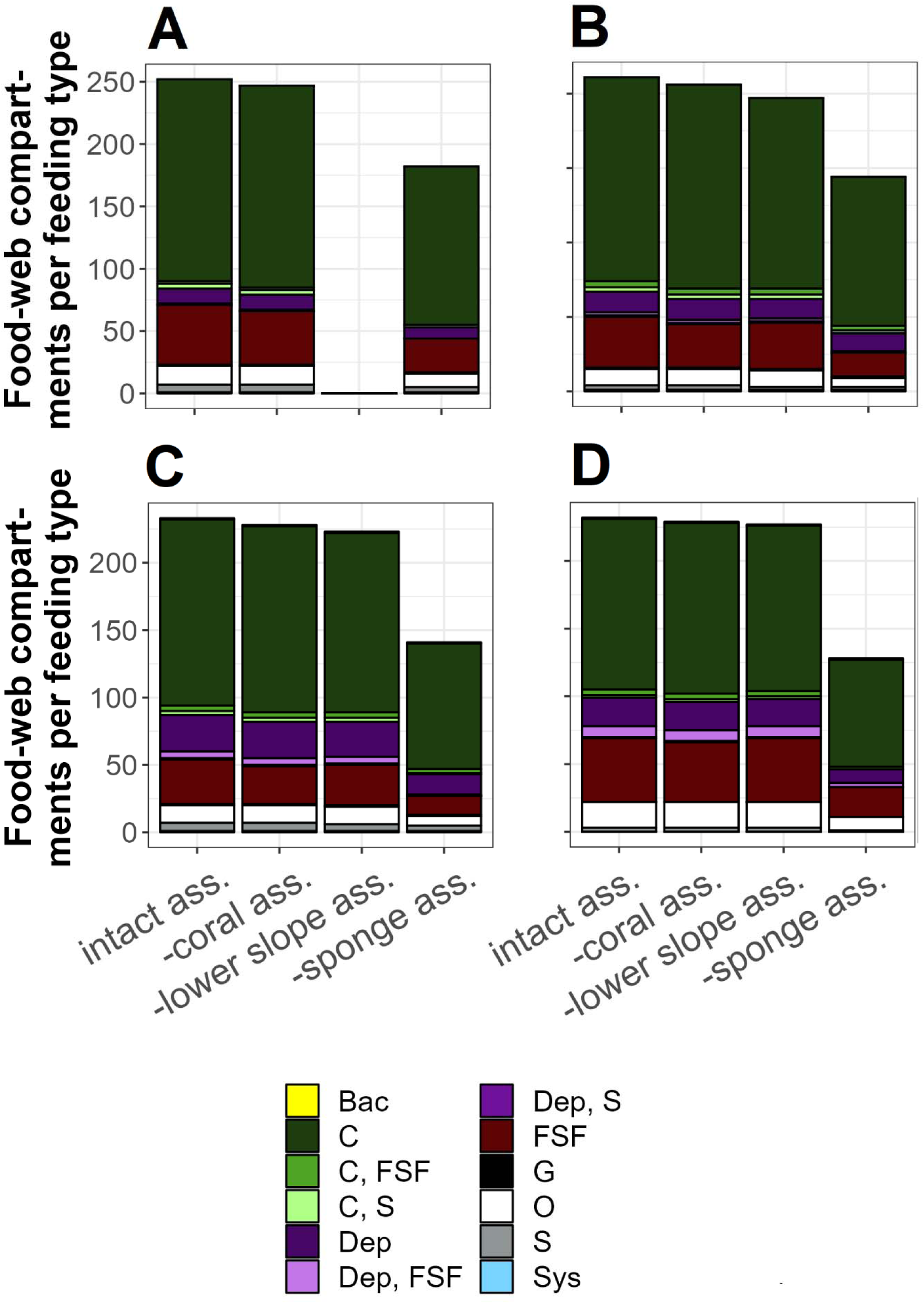
Number of feeding types present in the deep-sea coral assemblage (**A**), the lower slope assemblage 1 (**B**) and 2 (**C**), and the deep-sea sponge assemblage (**D**). All compartments that are part of trophic and non-trophic interaction webs are split into feeding types and shown as intact assemblage (intact ass.), in the absence of representatives of the deep-sea coral assemblage (-coral ass.), in the absence of representatives of the lower slope assemblages (-lower slope ass.), and in the absence of demosponges of the suborder Astrophorina (-sponge ass.). No data are shown for “-lower slope ass.” in panel (A) because no representative taxa of the lower slope assemblages were detected at sites of the deep-sea coral assemblage. Abbreviations: Bac = bacterivore, C = carnivore, Dep = deposit feeder, FSF = filter/ suspension feeder, G = grazer, O = omnivore, S = scavenger, Sys = symbiosis.

**Figure 4.**
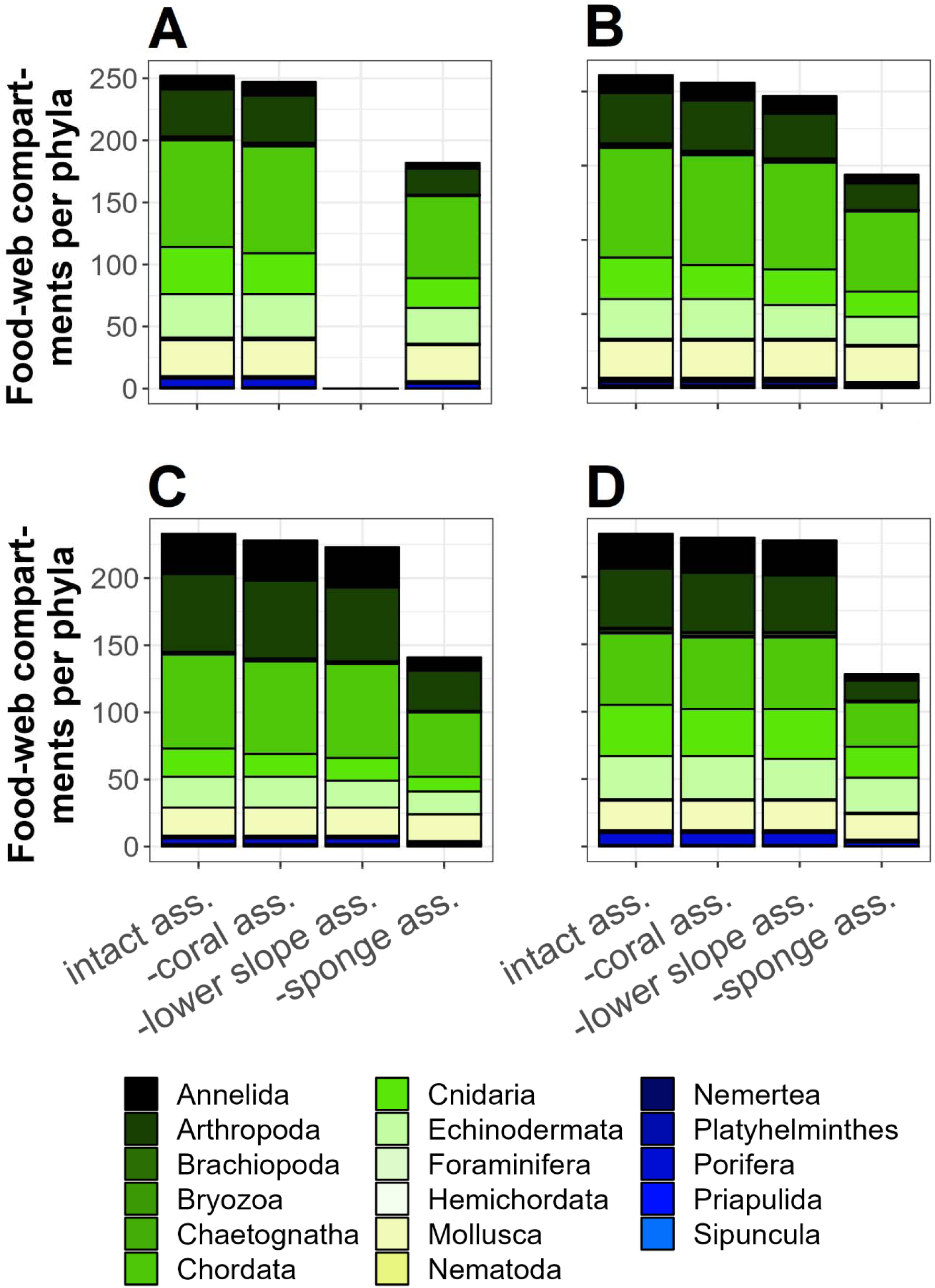
Number of phyla present in the deep-sea coral assemblage (**A**), the lower slope assemblage 1 (**B**) and 2 (**C**), and the deep-sea sponge assemblage (**D**). All compartments that are part of trophic and non-trophic interaction webs are divided into phyla and shown as intact assemblage (intact ass.), representatives of the deep-sea coral assemblage (-coral ass.), without representatives of the lower slope assemblages (-lower slope ass.), and without demosponges of the suborder Astrophorina (-sponge ass.). No data are shown for “-lower slope ass.” in panel (A) because no representative taxa of the lower slope assemblages were detected at sites of the deep-sea coral assemblage.

**Figure 5.**
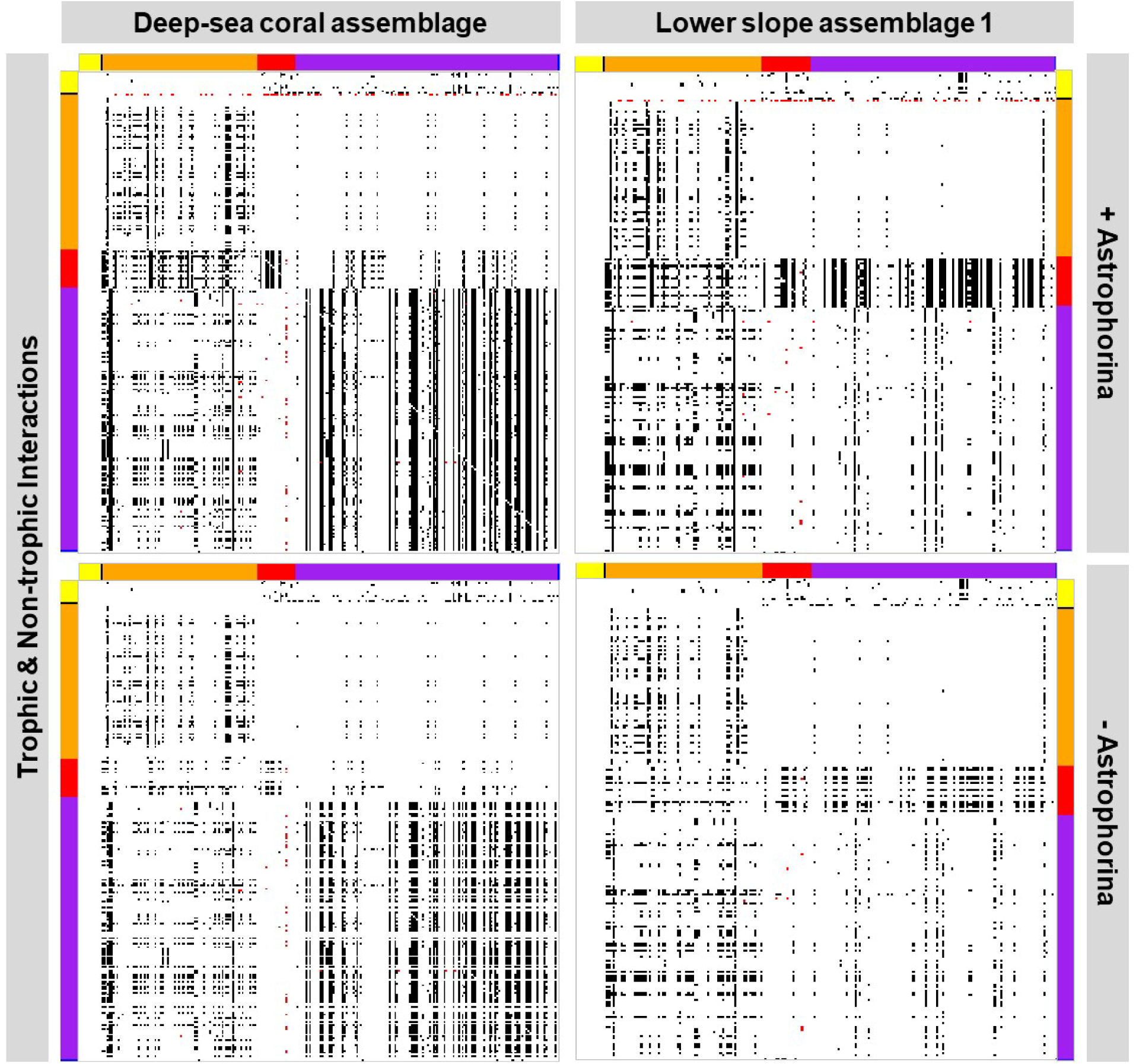
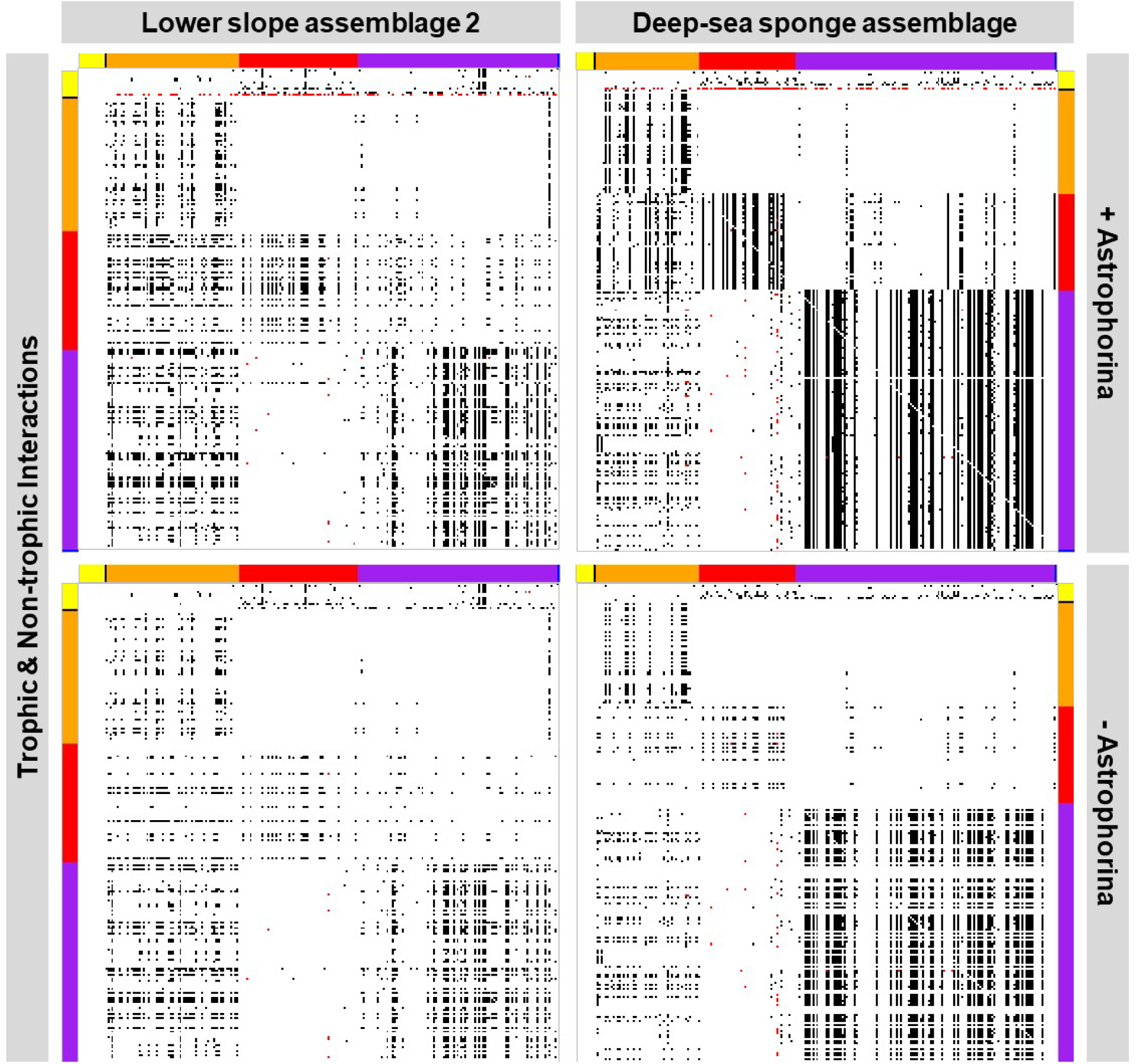
Network links between food-web compartments. Each pixel symbolizes an individual link between Astrophorina (black section) and another faunal food-web compartment (orange, red, purple, and blue sections), two different faunal food-web compartments, or a faunal food-web compartment and a food-source compartment (yellow section; Table 3). **(Top left)** Trophic (black) and non-trophic (red) links in the deep-water coral assemblage when sponges of the suborder Astrophorina are present. **(Bottom left)** All trophic (black) and non-trophic (red) links that remain in the deep-water coral assemblage interaction webs after Astrophorina removal. **(Top right)** All trophic (black) and non-trophic (red) links that occur in the lower slope assemblage 1 when Astrophorina are present. **(Bottom right)** All trophic (black) and non-trophic (red) links that remain after Astrophorina are removed in the lower slope interaction webs. Colour code of axes: yellow = food sources, black = sponges of the suborder Astrophorina, orange = fish, red = macrobenthos, purple = megabenthos, blue = meiobenthos. **(Top left)** All trophic (black) and non-trophic (red) links that occur in the lower slope assemblage 2 when Astrophorina are present. **(Bottom left)** All trophic (black) and non-trophic (red) links that remain after Astrophorina are removed in the cold-water coral interaction webs. **(Top right)** All trophic (black) and non-trophic (red) links that occur in the deep-sea sponge assemblage when Astrophorina are present. **(Bottom right)** All trophic (black) and non-trophic (red) links that remain after Astrophorina are removed in the deep-sea sponge interaction webs.

**Table 4.**
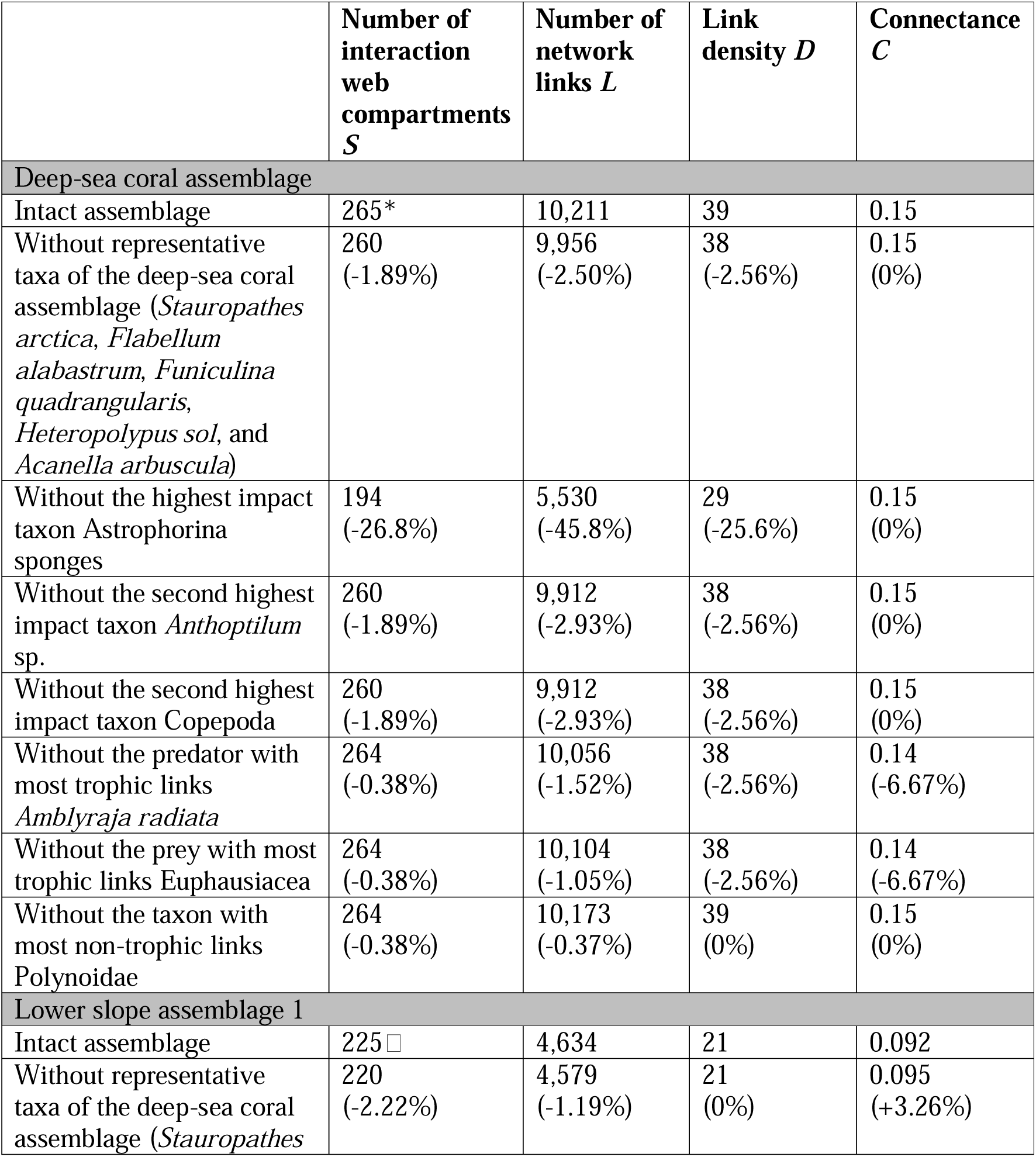
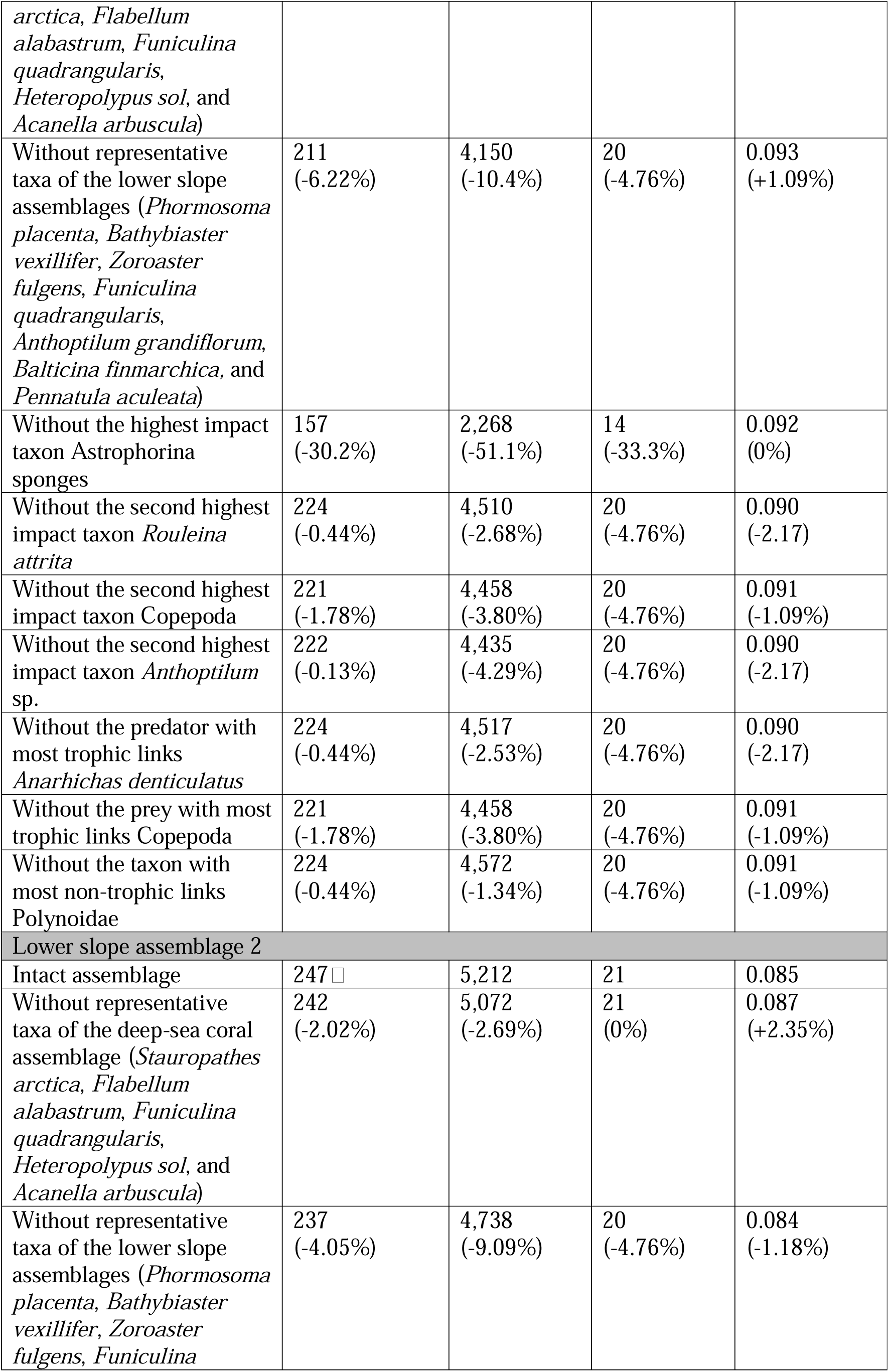
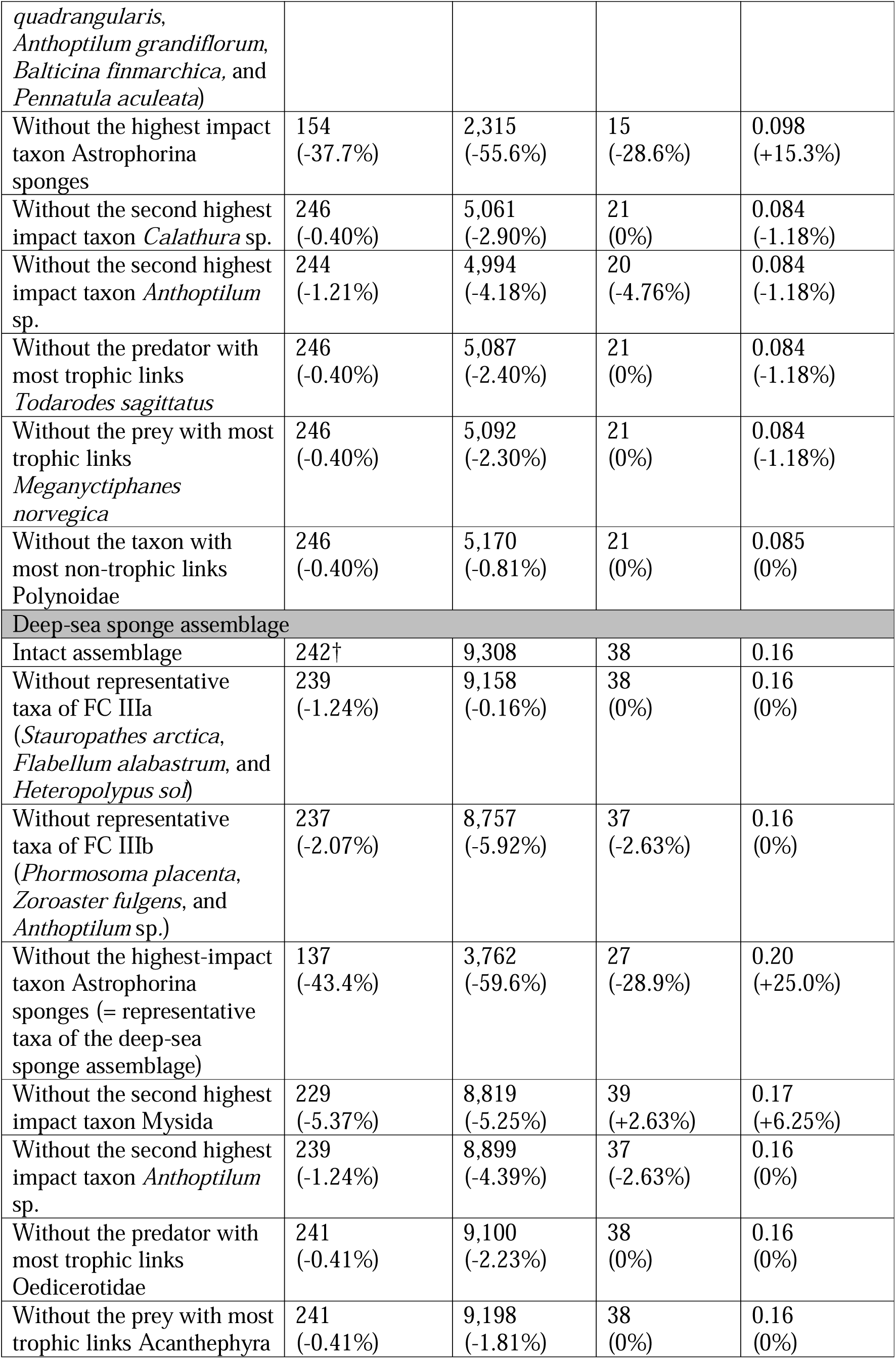
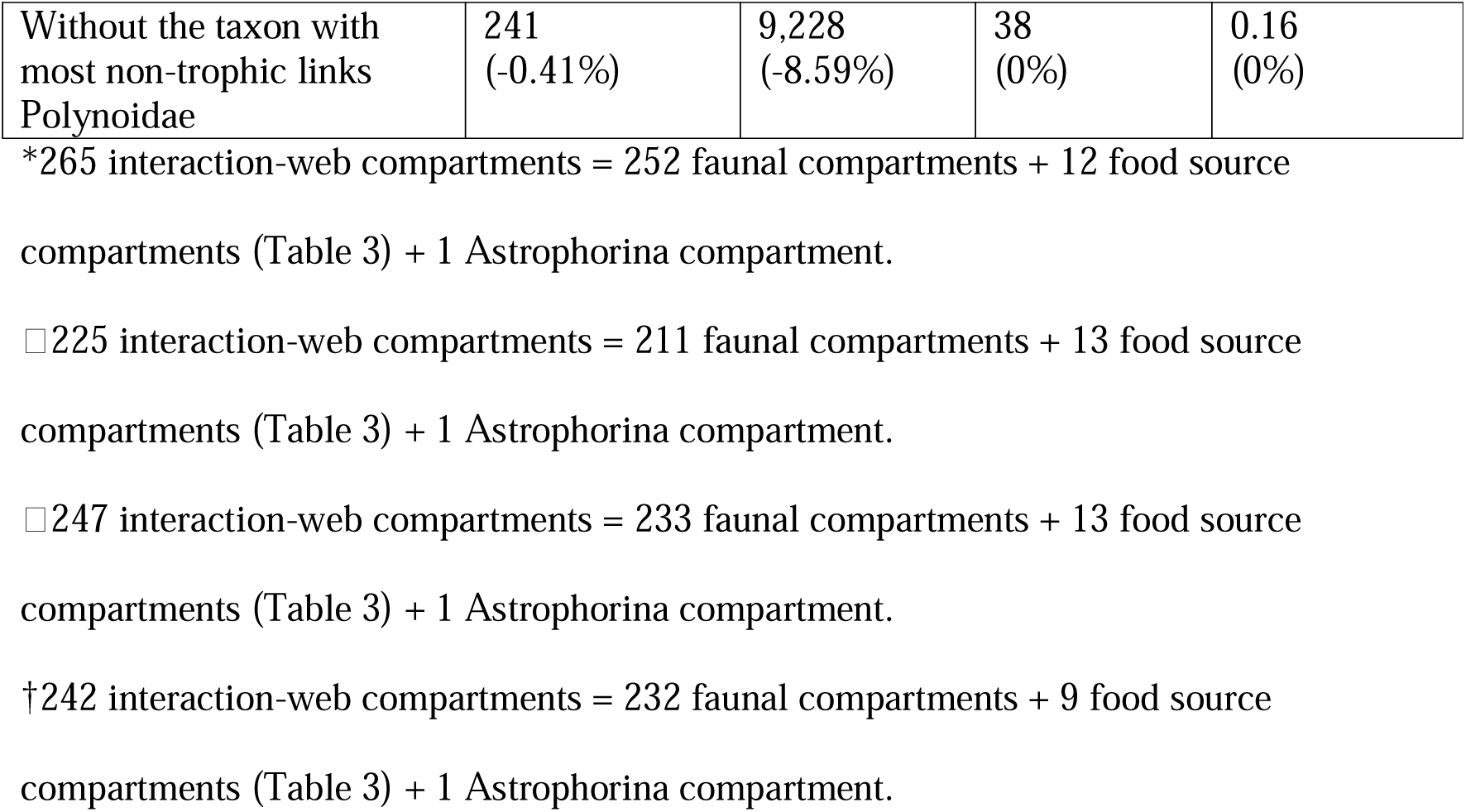
Changes of network properties dependent on the presence (intact assemblage) or absence of specific taxa. The network properties were calculated for the trophic interaction webs of the deep-sea coral assemblage, lower slope assemblages 1 and 2, and deep-sea sponge assemblage, and when the highest and second highest impact taxa were absent. The network properties were also calculated for the trophic interaction webs without the taxa with most trophic and non-trophic links. The changes in percent with respect to the default webs are shown in brackets.

#### 3.1.2 Lower slope assemblage 1

The lower slope assemblage 1 contained 211 faunal interaction web compartments of which 0.47% belonged to protozoan and metazoan meiobenthos, 10.9% to macrobenthos, 54.0% to invertebrate megabenthos, and 34.6% to fish. Most of the taxa were carnivores (64.9%) and filter/suspension feeders (16.1%) (Fig. 3B) and the dominant phyla were Chordata (35.1% of all taxa), Arthropoda (16.1%), Cnidaria (13.3%), Echinodermata (12.8%), and Mollusca (11.8%) (Fig. 4B). In total, 4,634 trophic links connected all compartments with their respective food sources (Fig. 5, Table 3, Table 4). The link density was 21 and the connectance was 0.092 (Table 4). Additionally, the compartments were connected with 162 links in the non-trophic interaction web (Fig. 5). The northern wolffish (*Anarhichas denticulatus*) was the predator with most trophic links, the prey with most trophic links was macrobenthic Copepoda, and polychaetes of the family Polynoidae had most non-trophic links.

#### 3.1.3 Lower slope assemblage 2

The lower slope assemblage 2 consisted of 233 faunal interaction-web compartments of which 0.86% were protozoan and metazoan meiobenthos, 26.2% were macrobenthos, 43.8% were invertebrate megabenthos, and 29.2% were fish. Most of the taxa were carnivores (59.5%), filter/suspension feeders (14.2%), and deposit feeders (11.6%) (Fig. 3C) and 67.8% of all faunal compartments were Annelida (12.9% of all taxa), Arthropoda (24.9% of all taxa), and Chordata (30.0% of all taxa) (Fig. 4C). All compartments were linked with 5,212 trophic links with their prey and other food sources (Fig. 5, Table 3, Table 4). Furthermore, 216 non-trophic links existed among compartments in the non-trophic interaction web (Fig. 5). The link density was 21 and the connectance was 0.085 (Table 4). The predator with most trophic links in the lower slope assemblage 2 was the megabenthic European flying squid (*Todarodes sagittatus*), the prey with most predators was Norwegian krill (*Meganyctiphanes norvegica*) and the taxon with most non-trophic links was the polychaete family Polynoidae.

#### 3.1.4 Deep-sea sponge assemblage

For the deep-sea sponge assemblage, 232 faunal interaction-web compartments were recorded ranging in size from protozoan and metazoan meiobenthos (0.86%), to macrobenthos (20.7%), invertebrate megabenthos (56.0%), and fish (22.4%). Seventy five percent of all compartments were carnivores (126 compartments), filter/ suspension feeders (47 compartments), and deposit feeders (21 compartments) (Fig. 3D). All faunal compartments were connected to their corresponding prey and food sources via 9,308 trophic links (Fig. 5, Table 3, Table 4). The non-trophic interaction web consisted of 318 links (Fig. 5). The link density was 38 and the connectance was 0.16 (Table 4). Members of the megabenthic amphipod family Oedicerotidae were the predators with the most trophic links, and shrimps (*Acanthephyra* sp.) were the prey with the most trophic links. Polychaetes of the family Polynoidae were the taxon with most non-trophic links.

### 3.2 Removing specific taxa from the different faunal assemblages

#### 3.2.1 Removing Astrophorina sponges

Removing the sponge suborder Astrophorina compartment from the deep-sea coral assemblage led to the loss of 26.8% of all interaction-web compartments (Table 4) due to first order removal of the facultative non-trophic (commensal) links with Astrophorina (Table 5), second order removal of facultative non-trophic links with other fauna, and maximum tertiary losses of trophic links. This indicates that Astrophorina sponges are the “highest impact taxa”. The lost compartments included macrobenthos (17.1% of all lost compartments), invertebrate megabenthos (55.7%), and fish (27.1%) (Fig. 5). Most of the removed faunal taxa were “carnivore, filter/ suspension feeder” (50.0%) and filter/ suspension feeder (43.8%). Additionally, the removal of Astrophorina resulted in the complete disappearance of the feeding types “carnivore and scavenger” and “deposit feeder and filter/ suspension feeder” (Fig. 3A). Phyla that disappeared due to the removal of Astrophorina were mostly Brachiopoda (100% loss), Bryozoa (100% loss), Sipuncula (100% loss), and Annelida (54.5% loss) (Fig. 4A).

**Table 5.**
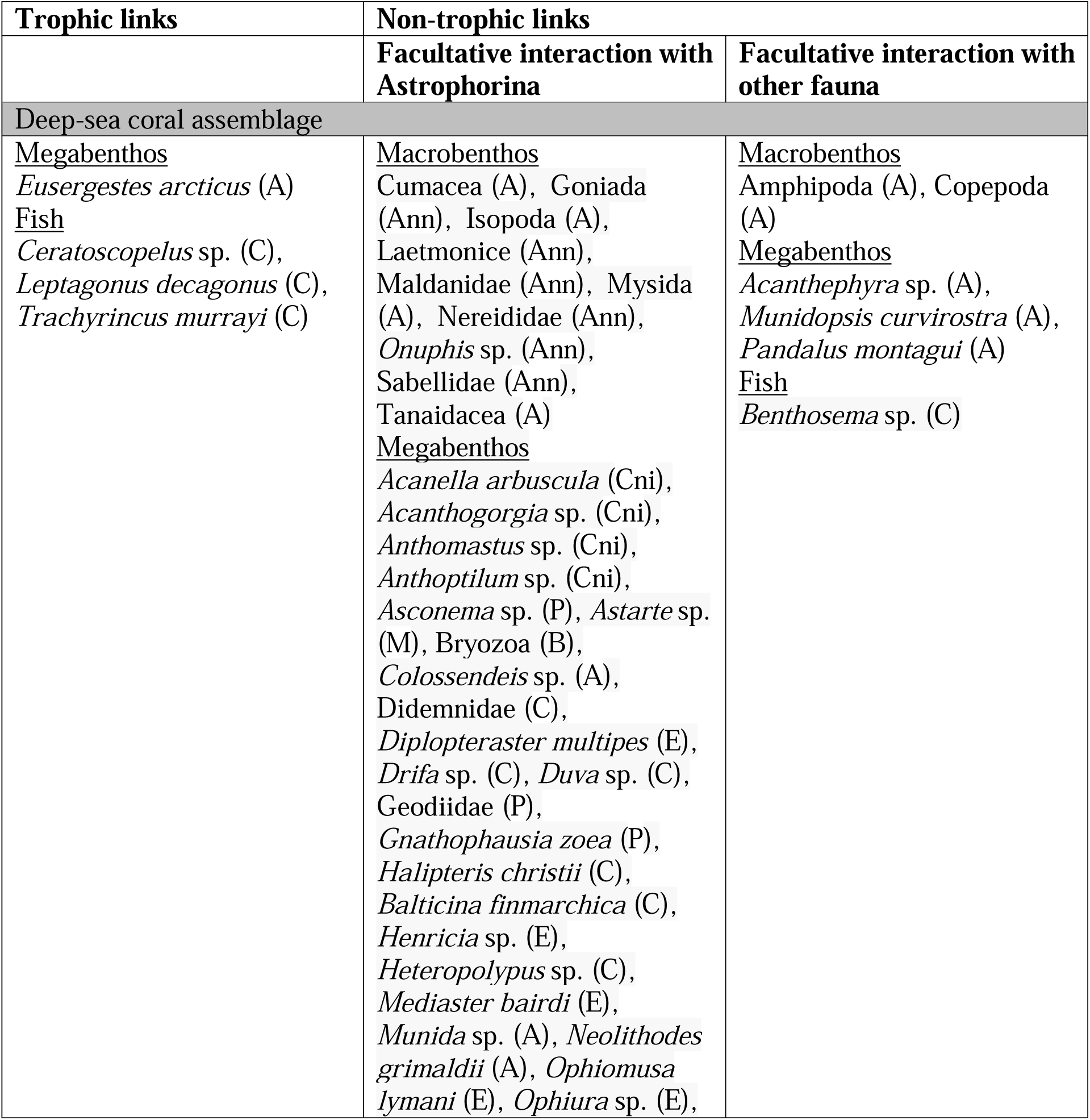
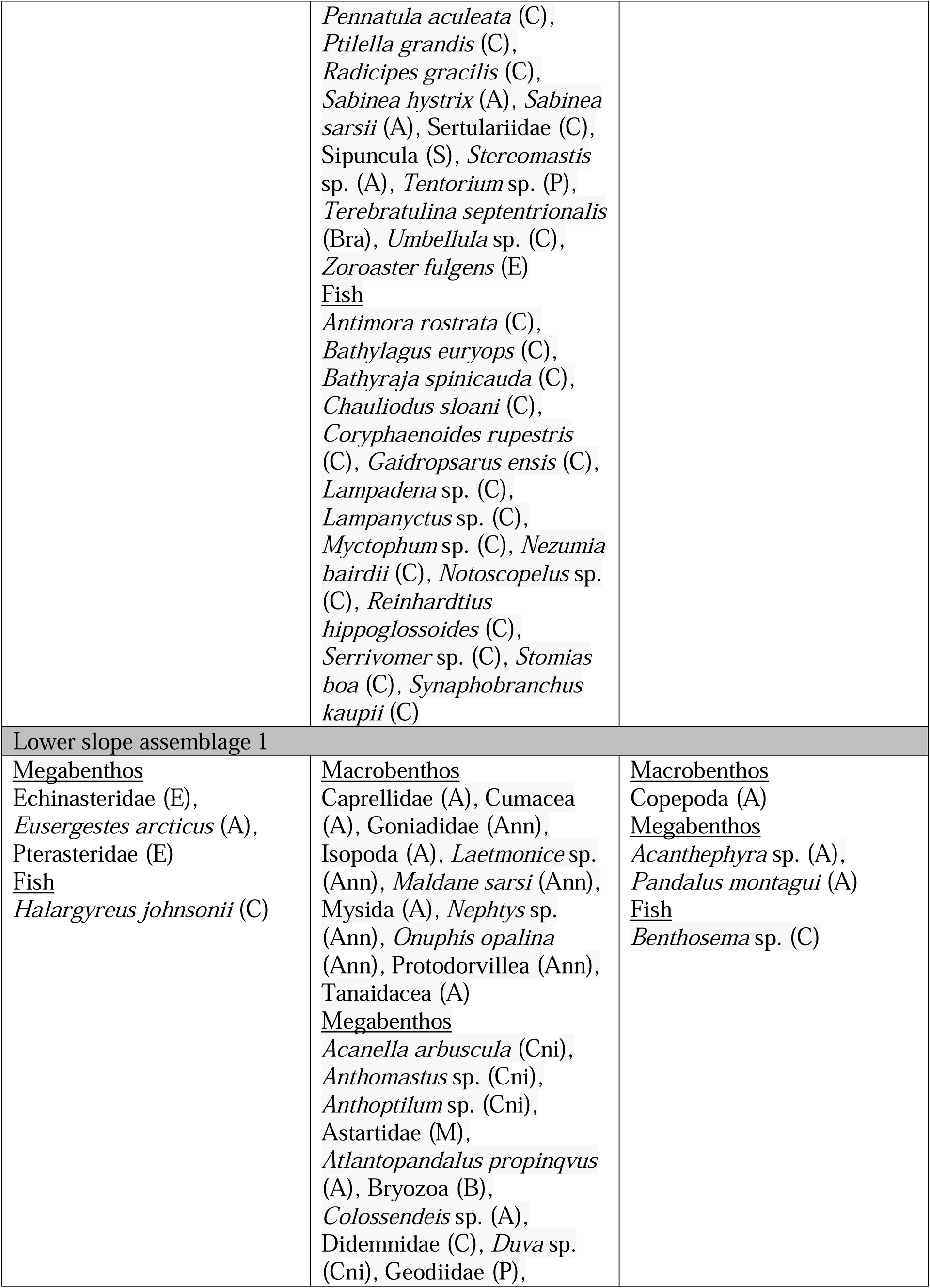
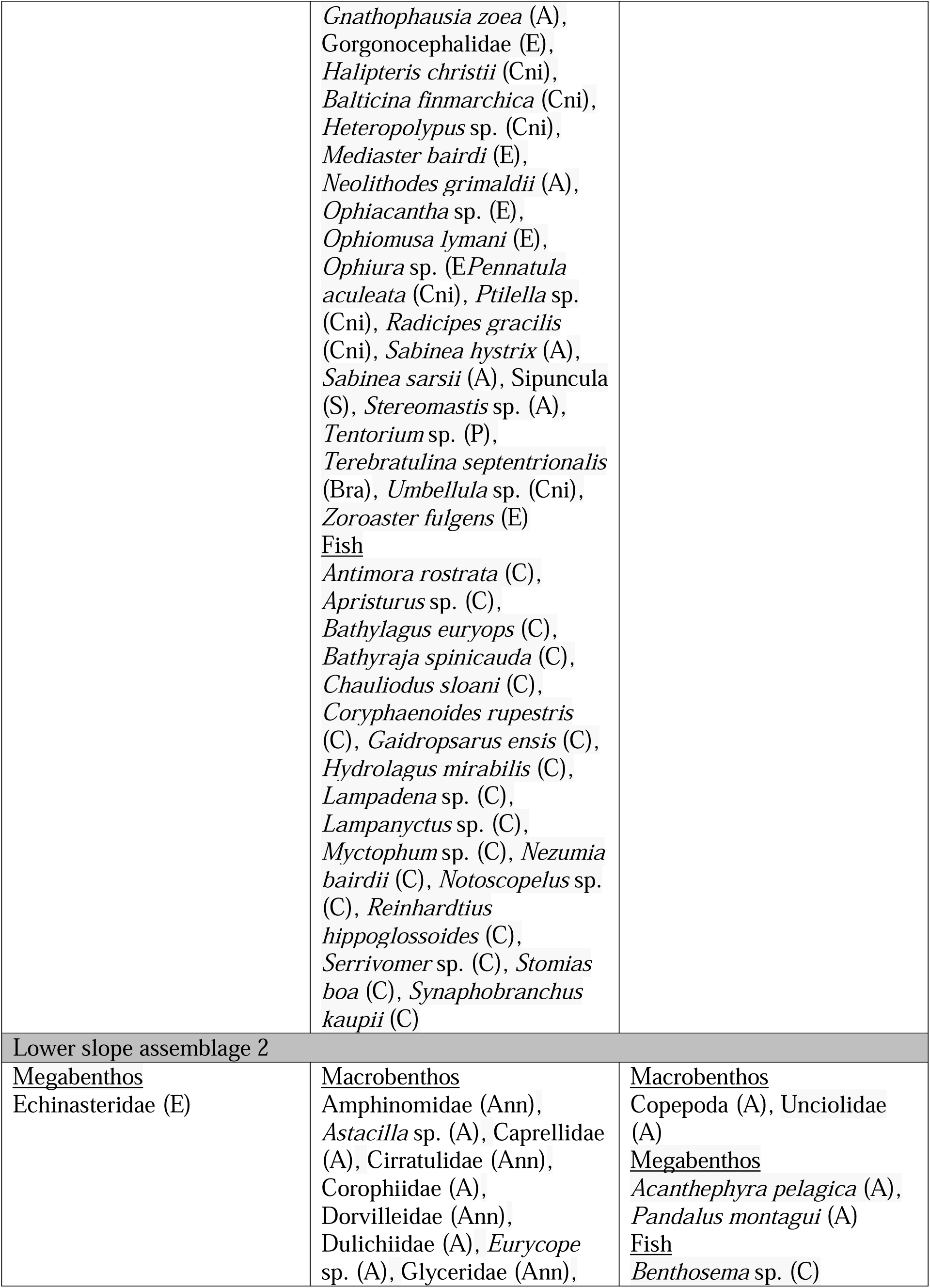
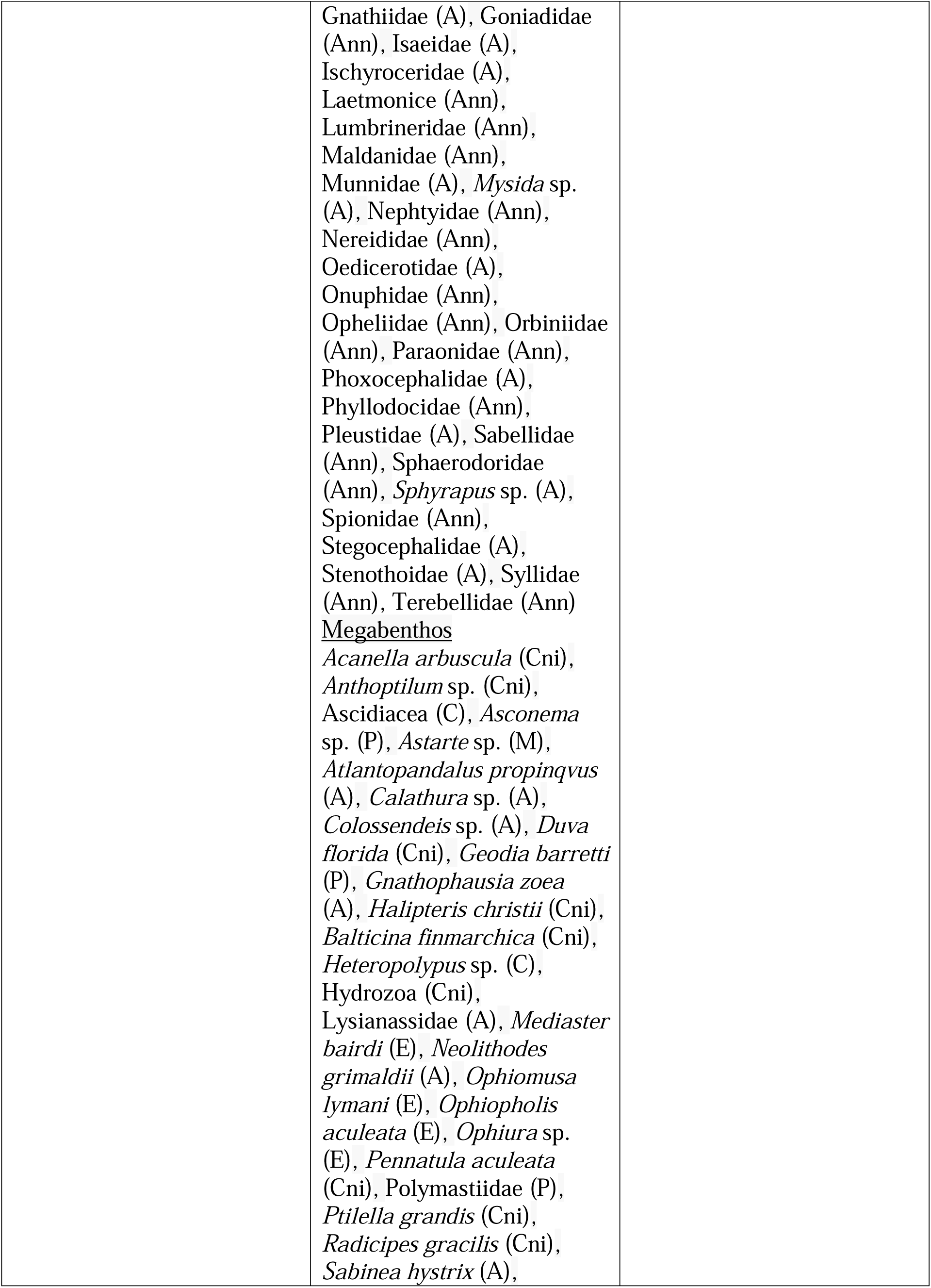
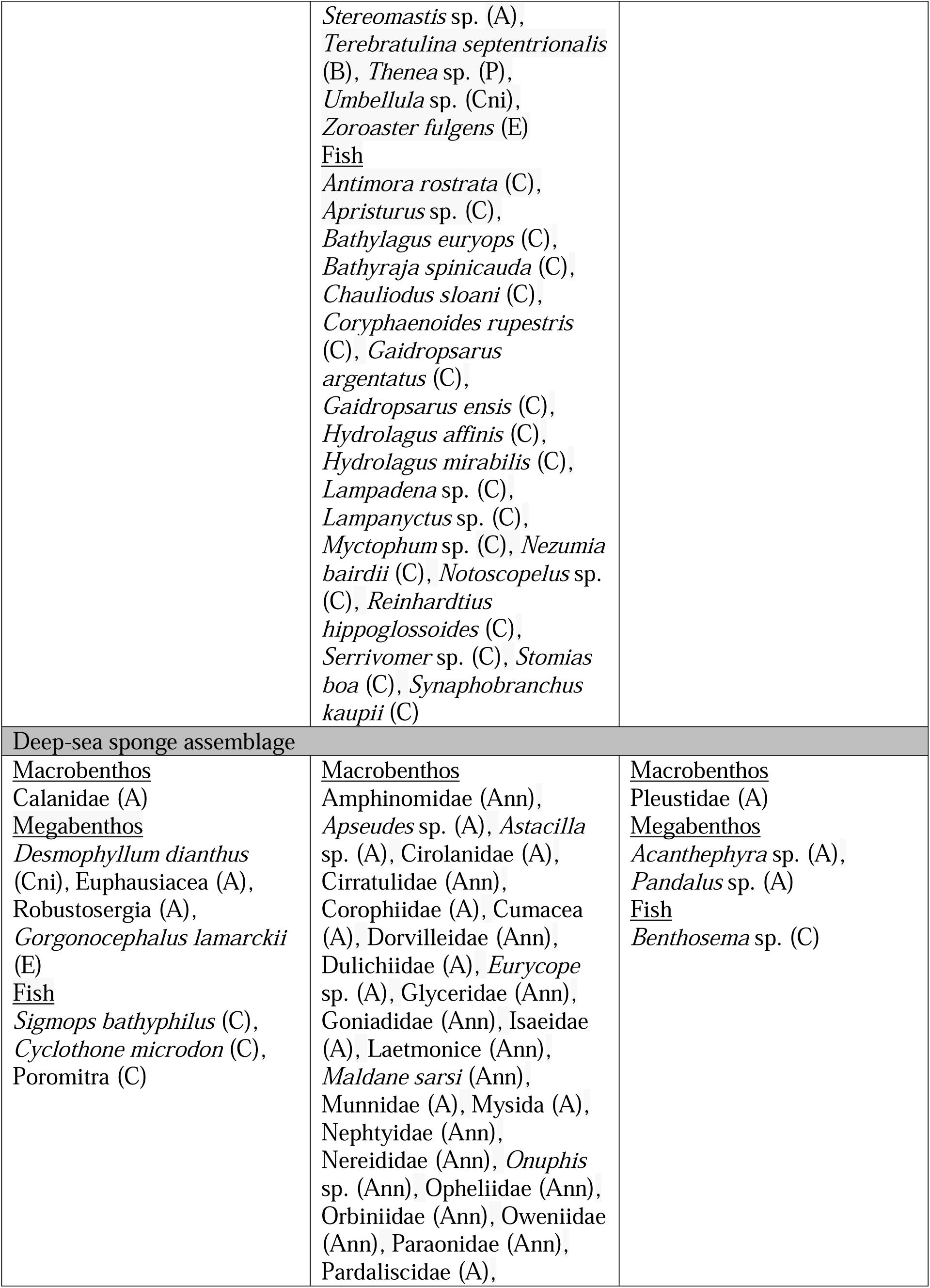
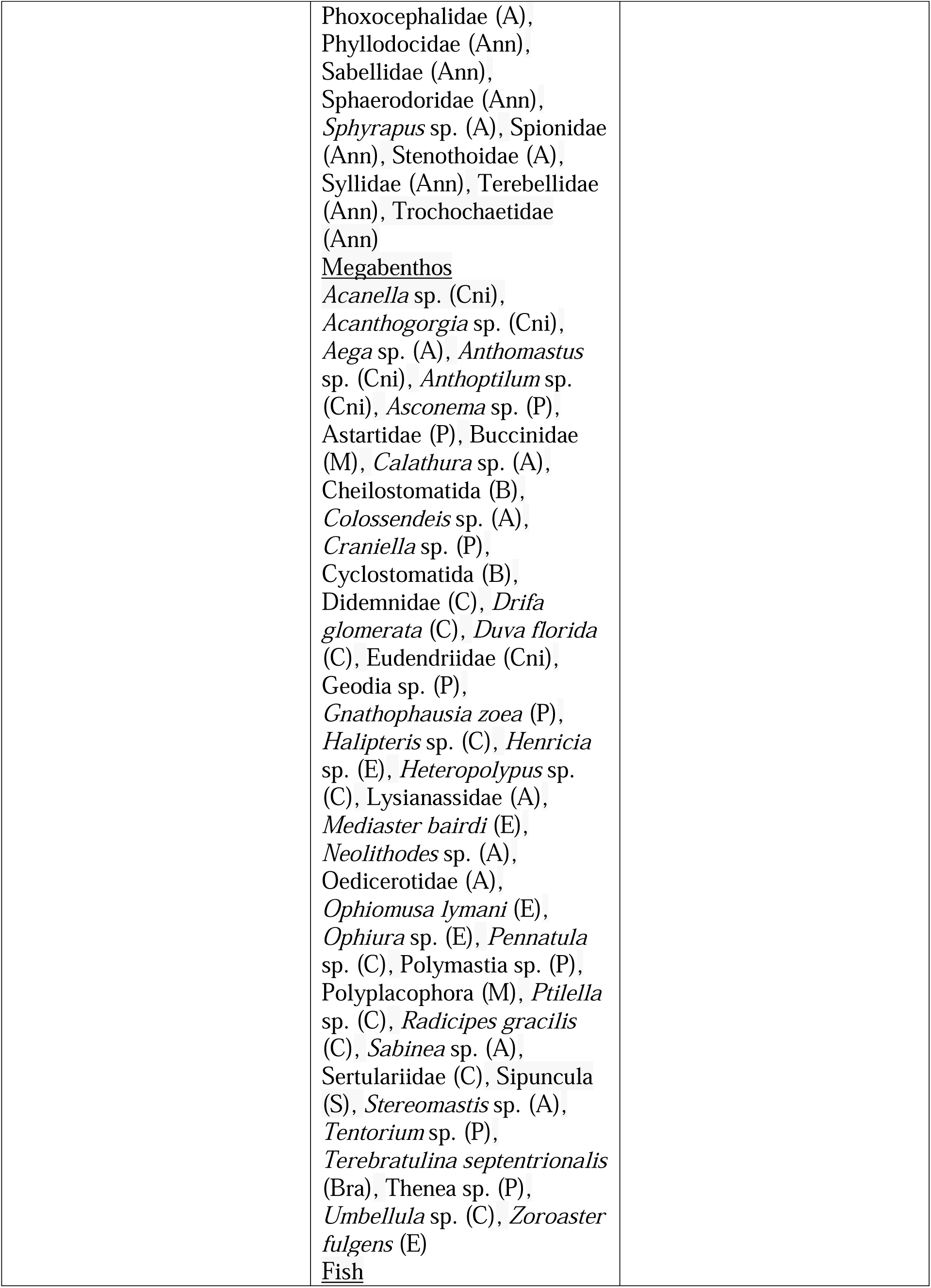
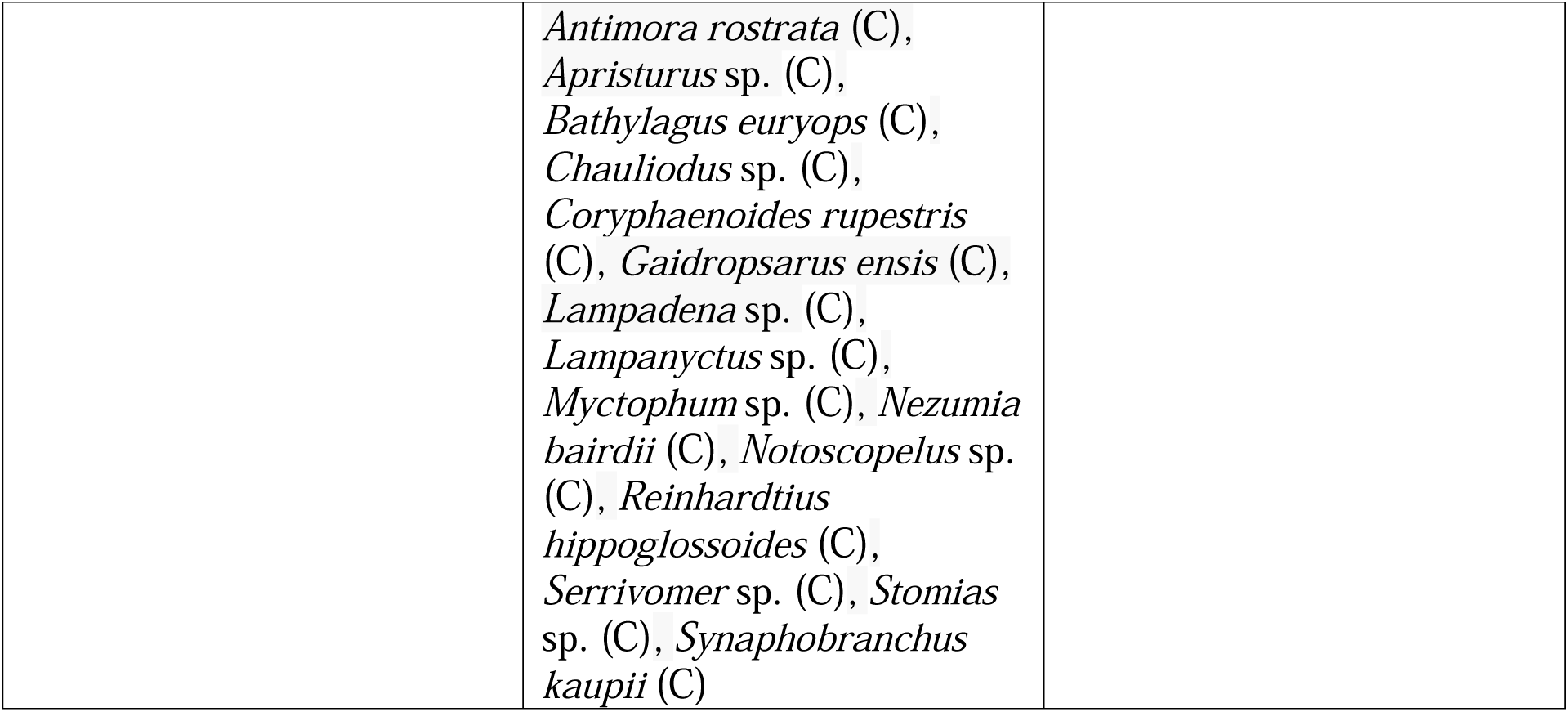
Faunal interaction web compartments lost by Astrophorina removal. All taxa are listed that disappear in the absence of Astrophorina due to trophic and non-trophic links \ with Astrophorina and other fauna. Letters in brackets behind the taxa indicate the phyla. Abbreviation of phyla: A = Arthropoda, Ann = Annelida, B = Bryozoa, Bra= Brachiopoda, C = Chordata, Cni = Cnidaria, E = Echinodermata, M = Mollusca, P = Porifera, S = Sipuncula.

The loss of the “highest impact taxon” Astrophorina from the lower slope assemblage 1 caused the removal of 67 taxa which was equivalent to the removal of 30.2% of all faunal compartments (Table 4) of which 17.9% belonged to macrobenthos, 53.7% to invertebrate megabenthos, and 28.4% to fish. Except for the megabenthic starfish families Echinasteridae and Pterasteridae, Arctic red prawn (*Eusergestes arcticus*) and slender codling (*Halargyreus johnsonii*) that disappeared from the interaction webs due to the loss of their food sources, all other lost faunal compartments had facultative non-trophic (commensal) links with Astrophorina or other fauna (Table 5). About 50% of taxa of the feeding types “deposit feeder and filter/ suspension feeder” and filter/ suspension feeder disappeared and the feeding type “deposit feeder and scavenger” was removed completely (Fig. 3B). Furthermore, all taxa of the phyla Brachiopoda, Bryozoa, Porifera, and Sipuncula were lost (Fig. 4B).

Eliminating the sponge suborder Astrophorina from the lower slope assemblage 2 resulted in the loss of 37.7% of all compartments (Table 4). Hence, Astrophorina sponges are the “highest impact taxa”. 41.3% of all macrobenthos, 37.0% of invertebrate megabenthos, and 21.7% of all fish disappeared mainly due to the loss of facultative links of taxa with Astrophorina sponges. Most of these lost taxa were carnivores (92 taxa), filter/ suspension feeder (19 taxa), and deposit feeders (12 taxa) (Fig. 3C), and they belonged mainly to the phyla Annelida (66.7% loss of all Annelida), Arthropoda (48.3% loss of all Arthropoda), Chordata (30.4% loss of all Chordata), and Cnidaria (50.0% of all Cnidaria) (Fig. 4C). Only megabenthic starfish of the family Echinasteridae was lost due to trophic links, whereas macrobenthic Copepoda and amphipods of the family Unciolidae were lost indirectly (i.e., maximum secondary loss) due to facultative links with the sea pen *Anthoptilum* sp., *Balticina finmarchica*, and *Umbellula* sp. The spiny pelagic deep-sea shrimp (*Acanthephyra pelagica*), the striped shrimp (*Pandalus montagui*), and the fish *Benthosema* sp. were facultatively non-trophically linked to the sea pen *Anthoptilum* sp. and were therefore lost in a second order loss.

Removal of the “highest impact taxon” sponge suborder Astrophorina from the deep-sea sponge assemblage resulted in a loss of 43.4% of all interaction-web compartments (Table 4), of which 35.7% belonged to the macrobenthic size class, 46.1% to invertebrate megabenthos, and 18.3% to fish. The removal of Astrophorina led to the disappearance of 37.3% of all carnivores, 47.4% of all omnivores, 52.4% of all deposit feeders, and 53.2% of all filter feeders (Fig. 3D). Macrobenthic Calanidae, megabenthic deep-water scleractinian coral *Desmophyllum dianthus*, the crustaceans Euphausiacea and Robustosergia, deep-sea basket star (*Gorgonocephalus lamarckii*), and the mesopelagic fish *Gonostoma bathyphilum*, veiled anglemouth (*Cyclothone microdon*), and Poromitra, were lost due to the loss of trophic links, i.e., due to the loss of their prey (Table 5). All other compartments were lost due to facultative non-trophic links with Astrophorina (88.5% of all compartment losses) and facultative links with other fauna (3.85% of all compartment losses). The last links contained the commensal relationship between the amphipod family Pleustidae and the octocoral *Acanthogorgia* sp. (though Pleustidae also had a facultative link to Astrophorina).

#### 3.2.2 Removing the representative taxa of the faunal assemblages

The disappearance of representative taxa of the deep-sea coral assemblage, i.e., the black coral *Stauropathes arctica*, the ivory stony coral (*Flabellum alabastrum*), the tall sea pen (*Funiculina quadrangularis*), the deep-sea mushroom soft coral (*Heteropolypus sol*), and the bonsai bamboo coral (*Acanella arbuscula*) from the assemblage did not result in the knock-down of any further compartments (Table 4). Removing representative taxa of the deep-sea coral assemblage from the lower slope assemblage 1 did not eliminate further compartments (Table 4). However, removing representative taxa of the lower slope assemblages, i.e., the flattened sea urchin (*Phormosoma placenta*), the seastars *Bathybiaster vexillifer* and *Zoroaster fulgens*, the tall sea pen, the full-flowered sea pen (*Anthoptilum grandiflorum*), *Balticina finmarchica,* and the thorny sea pen (*Pennatula aculeata*) from the lower slope assemblage 1 did eliminate compartments (i.e., macrobenthic Copepoda, *Acanthephyra* sp., striped shrimp, Acadian redfish *Sebastes fasciatus*) due to facultative non-trophic interactions with Astrophorina sponges and compartments (i.e., *Eusergestes arcticus*, Gorgonocephalidae, *Halargyreus johnsonii*) because of trophic interactions (Table 4). The removal of representative taxa of the deep-sea coral assemblage from the lower slope assemblage 2 did not cause a knock-down of any other faunal compartment (Table 4). However, removing representative taxa of the lower slope assemblages from the lower slope assemblage 2 led to the loss of macrobenthic Copepoda, the spiny pelagic deep-sea shrimp, and the striped shrimp. The loss of representatives of the deep-sea coral and lower slope assemblages caused the removal of the spiny pelagic deep-sea shrimp and the striped shrimp from the deep-sea sponge assemblage (Table 4).

#### 3.2.3 Removing taxa with most trophic/ non-trophic links and the second highest impact taxa

The removal of taxa with the most trophic and non-trophic links from the deep-sea coral assemblage interaction web did not result in the knock-down of any further compartments (Table 4). In comparison, the removal of the second-highest impact taxon (i.e., taxon whose removal resulted in the second largest changes in food-web properties) sea pen *Anthoptilum* sp. caused the disappearance of 1.89% of all compartments, as macrobenthic amphipods have facultative non-trophic links with the octocoral and larvae of the shrimps *Acanthephyra* sp. and the striped shrimp may use it as nursery (Baillon et al. 2014). In a maximum secondary loss, the lanternfish (*Ceratoscopelus* sp.) was lost due to the disappearance of its only prey, the amphipods. Removing the other second-highest impact taxon Copepoda resulted in the removal of Atlantic poache (*Leptagonus decagonus*), macrobenthic amphipods, and Arctic red prawn that all predate upon Copepoda, whereas the lanternfish was lost in a second order knockdown after the removal of its only prey from the interaction web, the amphipods.

Losing the northern wolffish as predator with the most prey and removing the taxon with most non-trophic links Polynoidea from the lower slope assemblage 1 did not result in the knock-down of any other compartments (Table 4). However, the removal of the prey taxon with most predators Copepoda from the interaction web resulted in the loss of slender codling (*Halargyreus johnsonii*), Arctic red prawn, and Gorgonocephalidae. Besides being the prey with most predators, Copepoda was also second highest impact taxon together with the sea pen *Anthoptilum* sp., and the soft-skin smooth-head (*Rouleina attrite*). The removal of the latter did not trigger any cascade of compartment losses, whereas removing the sea pen implied losing the crustaceans *Acanthephyra* sp. and striped shrimp.

Removing the taxa with the most trophic (i.e., Norwegian krill, European flying squid) and non-trophic (i.e., Polynoidae) links from the lower slope assemblage 2 had no further effect on any other faunal compartment (Table 4). Taxa, that had the second highest impact on food-web properties, were the sea pen *Anthoptilum* sp. and megabenthic *Calathura* sp. (Table 4), but only removing *Anthoptilum* sp. affected other interaction-web compartments, i.e., the striped shrimp and the spiny pelagic deep-sea shrimp.

The removal of the taxa with the most trophic (Oedicerotidae, Acanthephyra) and non-trophic links (Polynoidea) did not cause any knock-down of other compartments in the deep-sea sponge assemblage (Table 4). In comparison, removing one of the taxa with the second highest impact, the crustacean Mysida, caused the loss of 5.37% of all compartments (Fig. 5) in a three-level knock-down cascade. In this cascade, first, macrobenthic Calanidae disappeared due to the loss of their prey Mysida and in a second order loss, the fish *Sigmops bathyphilus* and veiled anglemouth, deep-water scleractinian coral *Desmophyllum dianthus*, the crustaceans Euphausiacea, *Robustosergia* sp., and deep-sea basket star were lost after the loss of their prey. Subsequently, in a third order loss, the lanternfishes (*Lampadena* sp., *Lampanyctus* sp., *Myctophum* sp.), the lancet fish (*Notoscopelus* sp.), and the ridgehead (*Poromitra* sp.) vanished.

### 3.3 Changes in network properties of different faunal assemblages at the Flemish Cap

Removal of the Astrophorina compartment in the deep-sea coral assemblage, the lower slope assemblages 1 and 2, and the deep-sea sponge assemblage led to a substantial decrease in the number of network links (deep-sea coral: 45.8%; lower slope 1: 51.1%; lower slope 2: 55.6%; deep-sea sponge: 59.6%) and link density (deep-sea coral: 25.6%, lower slope 1: 33.3%, lower slope 2: 28.6%, deep-sea sponge: 28.9%) (Fig. 5, Table 4). The connectance, however, was only affected by removing Astrophorina in the lower slope assemblage 2 (+15.3%) and in the deep-sea sponge assemblage (+25.0%), whereas the connectance of the deep-sea coral assemblage and lower slope assemblage 1 remained unaffected. Removing any of the taxa with most trophic and non-trophic links, or second highest impact taxa never changed the network indices by more than 9% (Table 4). The Polynoidae compartment removal in the deep-sea sponge assemblage, for example, caused the second largest change in the number of network links with a reduction by 8.59% (Table 4).

## 4. DISCUSSION

### 4.1 Data and model limitations

The highly resolved interaction webs of the four different faunal assemblages at the Flemish Cap were particularly strong for macrobenthos, invertebrate megabenthos, and demersal fish, but less for meiobenthos. This was related to the sampling gear deployed during previous research expeditions on which data this study was based: Bottom trawls are commonly used to collect invertebrate megabenthos and fish (Gage & Tyler 1991), though at the Porcupine Abyssal Plain (Northeast Atlantic), trawls underestimated invertebrate megabenthos biomass by a factor of 40 to ∼200 compared to photo surveys (Durden et al. 2017). USNEL boxcorers are appropriate to sample macrobenthos but are rarely used to collect meiobenthos samples (Danovaro 2010). Hence, we included only 3 different meiobenthos taxa on the order (Harpacticoida) and phylum levels (Nematoda, Foraminifera), though sponge grounds, cold-water coral gardens, and soft-sediment communities can have a high meiobenthos diversity (Raes & Vanreusel 2006, Sandulli et al. 2015). Therefore, we likely underestimated the complexity of the food webs of the four faunal assemblages. However, comparable trophic-/ non-trophic interaction web models developed for abyssal plains in the central Pacific and Southeast Pacific Ocean included 34% meiobenthos compartments, of which none were lost when the highest-impact taxa, i.e., the stalked sponges *Hyalonema* sp. and *Caulophacus* sp., were removed (Stratmann et al. 2021). Therefore, it could be expected that the results of our study are reasonably robust to the poorly resolved meiobenthos in our models.

It should be stressed that the consequences the removal of specific compartments has on the trophic/ non-trophic webs interaction webs constitute extreme cases as we assume that the links removed are the critical ones. While this is not an issue in relative terms (i.e., to assess which removals are generally expected to have a higher risk of impact), the results do not necessarily reflect true expected impacts.

### 4.2 Role of Astrophorina sponges in food webs at the Flemish Cap

In all four faunal assemblages, Astrophorina sponges were identified as “highest impact taxa” whose removal caused a loss of 27% (deep-sea corals) to 43% (deep-sea sponges) of all interaction-web compartments, 46% (deep-sea corals) to 60% (deep-sea sponges) of all links, 26% (cold-water corals) to 33% (lower slope 1) of the link density, and 0% (deep-sea corals) to 25% (deep-sea sponges) of connectance.

The deep-sea coral assemblage, whose representative taxa are corals and sea pens (Murillo et al. 2016b), is strongly linked with Astrophorina sponges via facultative, non-trophic links. In fact, these non-trophic links even include commensal links between the deep-sea mushroom soft coral, the bonsai bamboo coral and Astrophorina sponges. The bonsai bamboo coral has also been observed further north in submarine canyons off Newfoundland (Canada) where it forms large coral gardens with low species diversity (Baker et al. 2012). Though this species was found on fine sediments, it seems to prefer structures to anchor (Baker et al. 2012).

When all representative taxa of the deep-sea coral assemblage (*Stauropathes arctica*, *Flabellum alabastrum*, *Funiculina quadrangularis*, *Heteropolypus sol*, and *Acanella arbuscula*) were removed, only 2.5% of the links and 2.6% of the link density were lost, whereas the connectance remained unaffected (this study). This might be surprising at a first glance, because corals are usually considered (autogenic) “ecosystem engineers” (i.e., organisms that “change the environment via their own physical structure” and “modulate the distribution and abundance of other resources”; Jones et al., 1994), or “foundation species” (i.e., “organisms that provide structure, moderate local biotic and abiotic conditions, and have a large, positive effect on other spices in a community“; Angelini et al., 2011 *sensu* Dayton et al. (1974). However, the cold-water coral species present at the Flemish Cap are solitary species, including the scleractinian corals such as *Flabellum alabastrum*, and therefore do not provide three-dimensional coral reefs.

In contrast, the two sea pen species, *Anthoptilum grandiflorum* and *Balticina finmarchica*, that are both present at sites of the deep-sea coral assemblage, but not representative species of it (Murillo et al. 2016a), have been suggested to act as biogenic habitat (Baillon et al. 2014). Indeed, removing *Anthoptilum* sp. from the interaction web caused a two-order extinction cascade with the loss of four additional interaction-web compartments, that had facultative non-trophic interactions with the sea pen, and the maximum secondary loss of the lanternfish which disappeared due to the loss of its prey (Supplementary Material). In comparison, Astrophorina sponges at the deep-sea coral assemblage facultatively host up to 62 species which can be lost in two-order extinction cascades when these sponges are removed. Hence, the sea pens and Astrophorina sponges can both be considered “foundation species” as their links with other taxa are mainly of non-trophic nature (Ellison 2019) or “structural species”/ “habitat formers” since they control biodiversity like foundation species, but their impacts on ecosystem functions is unspecific (Ellison 2019). However, due to their possible role as a nursery area for fish and crustacean larvae (Baillon et al. 2012, 2014) and the low abundance of Astrophorina sponges on the soft sediment, the sea pens are likely more important for the deep-sea coral assemblage than the sponges.

The two lower slope assemblages 1 and 2 are represented by the flattened sea urchin, different starfishes, and sea pens. Starfishes of the families Echinasteridae and Pterasteridae predate upon Astrophorina sponges so that the removal of this sponge suborder from the interaction webs resulted in a loss of trophic links with these specific families. In fact, starfish are common spongivores: they predate upon *Geodia* sp. indet./ *Stelletta* sp. indet. at the Langseth Ridge in the central Arctic Ocean (Stratmann et al. 2021), upon various sponge species along the continental margin of the Northwest Atlantic Ocean and the Gulf of Mexico (Mah 2020), or upon reef sponges in the Caribbean (Wulff 1995). However, beyond being spongivores, the role of these starfishes in the food web of the Flemish Cap appears to be rather limited as their removal from the interaction webs of lower slope 1 and 2 did not affect any other taxon (Supplementary material).

Sea pens, in contrast, are often seen as biogenic habitat providers in soft-sediment habitats (Tissot et al. 2006, Baillon et al. 2014). At the continental margin off northern Norway (NE Atlantic), decapods of the families Caridea and Munididae were observed near 20% of the sea pens (De Clippele et al. 2015) and in the Laurentian Channel (Canada, NW Atlantic), macrobenthic diversity is (Miatta & Snelgrove 2022) higher at sites with sea pens compared to sites with bare sediment (Miatta & Snelgrove 2022). Like in the deep-sea coral assemblages, sea pens may provide nurseries for larvae of crustaceans and of the redfish (Baillon et al. 2012, 2014) in the lower slope assemblages and therefore, they have facultative non-trophic links with fish and arthropods, that are connected via trophic links to slender codling, Arctic red prawn, and sea urchins. Also, Astrophorina sponges have further facultative non-trophic links with a diverse range of annelids, cnidarians, arthropods, echinoderms, and fish in the lower slope assemblages.

The deep-sea sponge assemblage consists mainly of sponges of the suborder Astrophorina (i.e., *Geodia* sp., *Thenea* sp.) which can be considered “key species” (*sensu* Davic 2003) as they regulate energy and nutrient dynamics due to their large biomass (Murillo et al. 2012) and consequently, high rates of carbon and nitrogen cycling (Pham et al. 2019). They can also be considered “ecosystem engineers” because they module the distribution and abundance of resources (Jones et al. 1994), i.e., nutrients (Hoffmann et al. 2009). They are “structural species”/ “habitat formers” and create habitat for associated fauna (Ellison 2019). Since most of the links between Astrophorina sponges and other members of the interaction webs are non-trophic, they are “foundation species” (Ellison 2019) *sensu* Dayton (1972). In fact, Astrophorina sponges in the deep-sea sponge assemblage have facultative non-trophic links with 92 different taxa which are more trophic or non-trophic links than Astrophorina sponges have in any of the other investigated faunal assemblages.

Not only demosponges like the Astrophorina sponges have been identified as “foundation species”, but also glass sponges. Archer et al. (2020) developed food web models for 20 glass sponge reefs in British Columbia (Canada) and showed that these glass sponges are an important prey for all investigated taxa. This is very different from the role that Astrophorina sponges play at the Flemish Cap where they are mostly involved in non-trophic interactions. However, the diet information for the Flemish Cap is mostly taken from published literature, whereas Archer et al. (2020) based their food-web topology on stable isotope analysis and stomach content analysis from specimens collected at the sites for which they developed their models. Hence, we might underestimate the role of Astrophorina sponges in trophic links at the Flemish Cap due to a lack of site-specific diet information for potential predators. Interestingly, however, Archer et al. (2020) found out that the sponge cover has an influence on food-web topology: When the glass sponge cover is <10%, the food webs are less connected and first and second order consumers depend on fewer food sources and are predated upon by fewer predators. When the glass sponge coverage is >10%, the food web is more connected, and the species rely on more food sources but are also predated upon by more predators. In comparison, the faunal assemblage with the highest sponge coverage at the Flemish Cap, the deep-sea sponge assemblage, has only a 0.01 higher connectance than the faunal assemblage with the second-highest sponge coverage (deep-sea coral assemblage), but a 0.058 to 0.065 higher connectance than the lower slope assemblages 1 and 2 with the lowest sponge coverage. Hence, sponge coverage likely does not only affect connectance on a trophic level, but also on a non-trophic level in sponge-dominated ecosystems.

## 5. CONCLUSIONS

Sponges of the suborder Astrophorina play an important role in deep-sea coral, lower slope, and deep-sea sponge assemblages at the Flemish Cap. They can be considered “highest-impact taxa” for all faunal assemblages as their removal has the largest impact on food-web properties (i.e., number of interaction-web links, link density, connectance). They are also “structural species”/ “habitat formers” and “foundation species”, particularly for the deep-sea sponge community, where Astrophorina sponges additionally can be considered “ecosystem engineers”. Sea pens are second highest impact taxa in all assemblages and particularly important in the cold-water coral assemblage and the lower slope assemblages. These results can support the evaluation of the functional role of vulnerable marine ecosystems (VMEs) in the North Atlantic that the NAFO aims to protect.

## Acknowledgements

Data from the box corers and rock dredges used in this study was collected under the NAFO Potential Vulnerable Marine Ecosystems—Impacts of Deep-sea Fisheries project (NEREIDA). The project was supported by Spain’s General Secretary of the Sea (SGM), Spain’s Ministry for the Rural and Marine Environment, the Spanish Institute of Oceanography, the Geological Survey of Canada, the Canadian Hydrographic Service, the Ecosystem Research Division of Fisheries and Ocean Canada (DFO), the UK’s Centre for the Environment Fisheries and Aquaculture Science (Cefas), the Russian Polar Research Institute of Marine Fisheries and Oceanography, and the Russian P.P. Shirshov Institute of Oceanology (RAS). The authors would like to acknowledge the hard work of the crew and scientists aboard the Spanish RV *Miguel Oliver* who collected the samples analyzed in this study. Groundfish surveys were co-funded by the EU through the European Maritime and Fisheries Fund (EMFF) within the National Program of collection, management and use of data in the fisheries sector and support for scientific advice regarding the Common Fisheries Policy. We further thank Cam Lirette (DFO) for his assistance in compiling the data for our use and our colleagues in the Northwest Atlantic Fisheries Organization (NAFO) Working Group on Ecosystem Science and Assessment (WG-ESA).

TS was supported by the Dutch Research Council NWO (NWO-Rubicon grant no. 019.182EN.012, NWO-Talent program Veni grant no. VI.Veni.212.211).

## Data

The R.file with the functions, the input files the interaction web models, and the Rmarkdown files with the results are published here: 10.5281/zenodo.7902500

